# Astrocyte transcriptomic changes along the spatiotemporal progression of Alzheimer’s disease

**DOI:** 10.1101/2022.12.03.518999

**Authors:** Alberto Serrano-Pozo, Zhaozhi Li, Maya E. Woodbury, Clara Muñoz-Castro, Astrid Wachter, Rojashree Jayakumar, Annie G. Bryant, Ayush Noori, Lindsay A. Welikovitch, Miwei Hu, Karen Zhao, Fan Liao, Gen Lin, Timothy Pastika, Joseph Tamm, Aicha Abdourahman, Taekyung Kwon, Rachel E. Bennett, Robert V. Talanian, Knut Biber, Eric H. Karran, Bradley T. Hyman, Sudeshna Das

**Author notes:** Equal contribution (Alberto Serrano-Pozo, Zhaozhi Li). **Correspondence to:** Sudeshna Das, PhD 65 Landsdowne St., Cambridge, MA 02139.

## Abstract

Astrocytes play a critical role in brain homeostasis and normal functions but their changes along the spatiotemporal progression of Alzheimer’s disease (AD) neuropathology remain largely unknown. Here we performed single-nucleus RNA-sequencing on brain regions along the stereotypical progression of AD pathology from donors ranging the entire normal aging-AD continuum comprising 628,943 astrocyte nuclei from 32 donors across 5 brain regions. We discovered temporal gene-expression-trajectories with gene sets differentially activated at various disease stages. Surprisingly, a gene set enriched in proteostasis and energy metabolism, was upregulated in late-stage but unexpectedly returned to baseline levels in end-stage, suggesting exhaustion of response in “burnt-out” astrocytes. The spatial gene-expression-trajectories revealed that astrocytic genes of tripartite synapses are dysregulated in parallel to the stereotypical progression of tangle pathology across regions. We identified astrocyte heterogeneity across brain regions with a continuum from homeostatic to reactive cells through “intermediate” transitional states. These findings suggest complex astrocytic dysfunction in AD neurodegeneration.

## INTRODUCTION

Alzheimer’s disease (AD) is defined by widespread accumulation of amyloid-*β* (A*β*) plaques and phospho-tau (pTau) neurofibrillary tangles (NFTs) throughout the brain, with the latter following a stereotypical hierarchical spatiotemporal pattern along neural networks. These AD neuropathological changes (ADNC) are accompanied by a dramatic loss of synapses and neurons as well as prominent morphological and functional changes of astrocytes, collectively termed reactive astrogliosis. Astrocytes are critical for maintaining brain homeostasis,^1^ and the view emerging from experimental data is that reactive astrogliosis may contribute to neurodegeneration through both gains of toxic functions and losses of normal functions^2^. Recent single-nucleus RNA-sequencing (snRNA-seq) studies^3–9^ have begun to unravel the molecular underpinnings of AD reactive astrocytes, but several questions remain: (1) Are there regional differences in astrocyte gene expression in the normal aging brain? (2) What are the gene expression changes that astrocytes exhibit along the spatiotemporal progression of AD and how does the severity of ADNC affect the astrocyte transcriptome? (3) Are there different astrocyte transcriptional subpopulations or states in the AD brain? We addressed these questions by conducting a large snRNA-seq study in five brain regions from 32 control and AD donors with various degrees of ADNC.

## RESULTS

### A molecular survey of astrocytes in five brain regions affected in a stereotypical fashion in AD

Using an enrichment strategy consisting of depleting NEUN+ve neurons and OLIG2+ve oligodendrocytes via fluorescence-activated nuclei sorting (FANS)^8^ we obtained a total of 628,943 astrocyte nuclei from five brain regions of 32 individuals with autopsy findings along the normal aging-AD neuropathological continuum (Figure 1a). The five brain regions were selected to represent the hierarchical spreading of pTau NFTs along brain networks as categorized by Braak NFT staging^10, 11^, and included entorhinal cortex (EC), inferior temporal gyrus (ITG, Brodmann Area [BA] 20), dorsolateral prefrontal cortex (PFC, BA46), visual association cortex (V2, BA18/19), and primary visual cortex (V1, BA17). Our strategy to enrich astrocyte nuclei was effective as indicated by the low numbers of neuronal and oligodendroglial nuclei identified (Figure 1b). Astrocyte nuclei were identified by the expression levels of marker genes *ADGRV1*, *ALDH1L1*, *AQP4*, and *GFAP* (Figure 1c), resulting in over 100,000 astrocyte nuclei in each brain region with *≍*6,000 average total number of reads and *≍*2,500 average total number of genes detected, which is orders of magnitude higher than previous studies^3–9^ (Supplementary Figure 1a).

**Figure 1.**
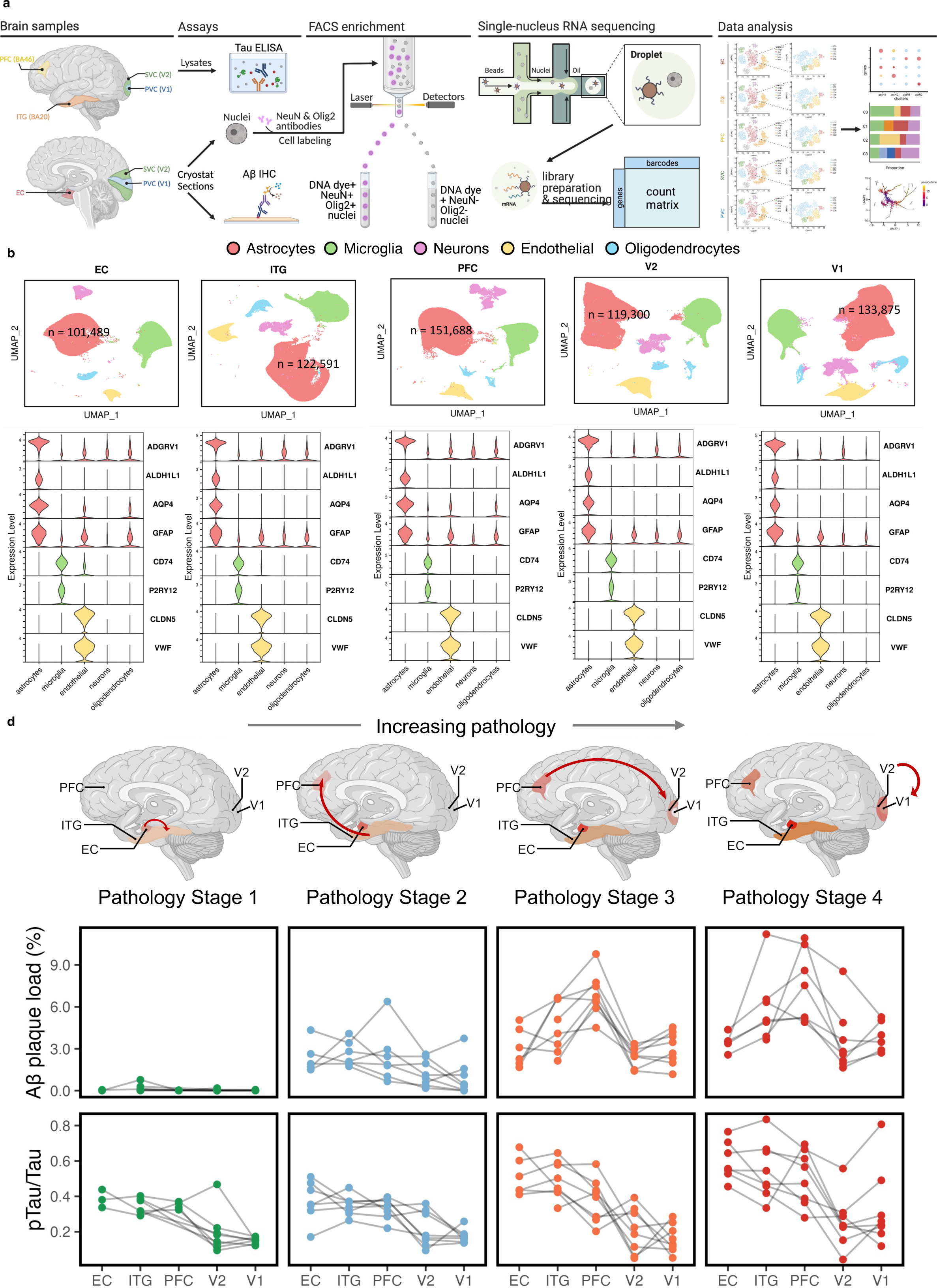
A molecular survey of astrocytes in five brain regions affected in a stereotypical manner in AD. **a**, Experimental overview of our snRNA-seq study. **b**, UMAP visualization showing clustering of NEUN-/OLIG2-nuclei. **c**, Violin plots illustrate the expression levels of cell type-specific markers genes in NEUN-/OLIG2-nuclei across the five brain regions. **d**, Results of Aβ plaque load (% immunoreactive area fraction) and pTau/Tau ratio (measured by ELISA) in adjacent samples to those used for snRNA-seq across brain regions and pathology stages. EC = entorhinal cortex; ITG = inferior temporal gyrus (BA20); PFC = prefrontal cortex (BA46); V2 = secondary (association) visual cortex (BA18/19); and V1 = primary visual cortex (BA17).

Since astrocytes react to nearby A*β* plaques and pTau NFTs^12, 13^, we quantified the local burden of ADNC in adjacent tissue samples by measuring the 3D6+ve A*β*-immunoreactive % area fraction via immunohistochemistry and the pTau/total-tau ratio via ELISA. To reflect the progression of ADNC, we then grouped the 32 donors into four pathology stages based on their global semiquantitative measures of neuritic plaques (CERAD NP score) and NFTs (Braak NFT stages) complemented with these immunohistochemical and biochemical quantitative measures of local A*β* and pTau burdens. The four pathology stages were: (1) Not AD/low ADNC burden (no NPs, Braak NFT stage 0/I/II); (2) intermediate ADNC burden (sparse or moderate NPs and Braak NFT stage II/III); (3) high ADNC burden with moderate or frequent NPs and Braak NFT stage V; and (4) high ADNC burden with moderate or frequent NPs and Braak NFT stage VI. We separated the latter two groups because, by definition, the primary visual cortex (region V1) bears NFTs only in Braak NFT stage VI^10, 11^. Figure 1d illustrates the quantitative measures of ADNC across brain regions of the 32 donors grouped in these four pathological stages. Within each stage, the A*β* plaque load was relatively constant across brain regions except for higher levels in PFC in late stages. By contrast, the pTau/Tau ratio was highest in EC and demonstrated the expected pattern EC>ITG>PFC>V2>V1 in all stages, consistent with the stereotypical hierarchical accumulation of NFTs along neural networks^10, 11^ (Figure 1d and Supplementary Figure 1b). The demographic and neuropathological characteristics of the study donors, including quantitative measures of A*β* and pTau, are summarized in Table 1 and detailed in Supplementary Table 1.

**Table 1:** Donor Characteristics

| Summary | Pathology Stage 1 | Pathology Stage 2 | Pathology Stage 3 | Pathology Stage 4 |
| --- | --- | --- | --- | --- |
| Age at death (mean ± SD) | 79.5 ± 12.3 | 86 ± 6.6 | 82.875 ± 7.9 | 79.5 ± 10.4 |
| Sex, N (% Female) | 4 (50) | 5 (62.5) | 6 (75) | 4 (50) |
| APOE ε4 carriers, N (%) | 0 (0) | 0 (0) | 3 (37.5) | 4 (50) |
| Braak NFT stage | 0/I/II | II/III | V | VI |
| CERAD NP score | none | sparse or moderate | moderate or frequent | moderate or frequent |
| Aβ plaque burden (mean ± SD) | 0.070 ± 0.152 | 1.855 ± 1.384 | 4.072 ± 2.103 | 4.909 ± 2.536 |
| EC | 0.031 ± 0.036 | 2.420 ± 1.153 | 3.072 ± 1.380 | 3.484 ± 0.571 |
| BA20 | 0.192 ± 0.274 | 2.513 ± 0.845 | 4.662 ± 1.842 | 5.938 ± 2.568 |
| BA46 | 0.018 ± 0.010 | 2.448 ± 1.900 | 6.869 ± 1.539 | 7.556 ± 2.544 |
| V2 | 0.040 ± 0.061 | 1.271 ± 0.943 | 2.467 ± 0.701 | 3.524 ± 2.278 |
| V1 | 0.021 ± 0.019 | 0.908 ± 1.277 | 3.039 ± 1.180 | 3.894 ± 1.072 |
| pTau/Tau (mean ± SD) | 0.257 ± 0.114 | 0.286 ± 0.109 | 0.343 ± 0.178 | 0.443 ± 0.204 |
| EC | 0.385 ± 0.051 | 0.374 ± 0.114 | 0.512 ± 0.107 | 0.600 ± 0.109 |
| BA20 | 0.343 ± 0.043 | 0.347 ± 0.056 | 0.494 ± 0.104 | 0.547 ± 0.162 |
| BA46 | 0.348 ± 0.021 | 0.333 ± 0.058 | 0.383 ± 0.124 | 0.505 ± 0.152 |
| V2 | 0.193 ± 0.119 | 0.212 ± 0.100 | 0.208 ± 0.128 | 0.256 ± 0.147 |
| V1 | 0.150 ± 0.017 | 0.175 ± 0.038 | 0.162 ± 0.079 | 0.328 ± 0.221 |

### Regional heterogeneity of astrocyte transcriptome in the normal aging brain

First, we asked whether astrocyte transcriptomes vary across regions in the normal aging brain. The eight control donors provided a unique opportunity to examine regional differences in the astrocyte transcriptome in the normal aging brain because these donors had essentially no 3D6+ve A*β* plaques across the five brain regions and low pTau/Tau ratios except for the EC, as expected in Braak NFT stages 0/I/II (Figure 1d). To address this question, we integrated a total of 246,464 nuclei from all brain regions of these eight individuals and conducted a differential gene expression analysis of each brain region relative to all other regions (Supplementary Table 2). Remarkably, the EC and the V1 regions had the highest number of differentially expressed genes (DEGs) (EC: 137 up and 152 down; V1: 93 up and 122 down, see Venn diagrams in Figure 2a). A heatmap with relevant DEGs per region is shown in Figure 2b. Among other genes of interest, *APOE*, *APP*, *AQP4*, and *GJA1* were upregulated, whereas *ALDH1L1*, *LRP1B*, *MAPT*, and *SLC1A2* were downregulated in EC relative to the other brain regions. Conversely, *CLU*, *GFAP*, and *MAOB* were upregulated, whereas *AQP4* and *GJA1* were downregulated in V1 vs. all other brain regions. Validation with immunohistochemistry confirmed regional differences between EC and V1 for *AQP4* and *GJA1* at the protein level (Figure 2c).

**Figure 2.**
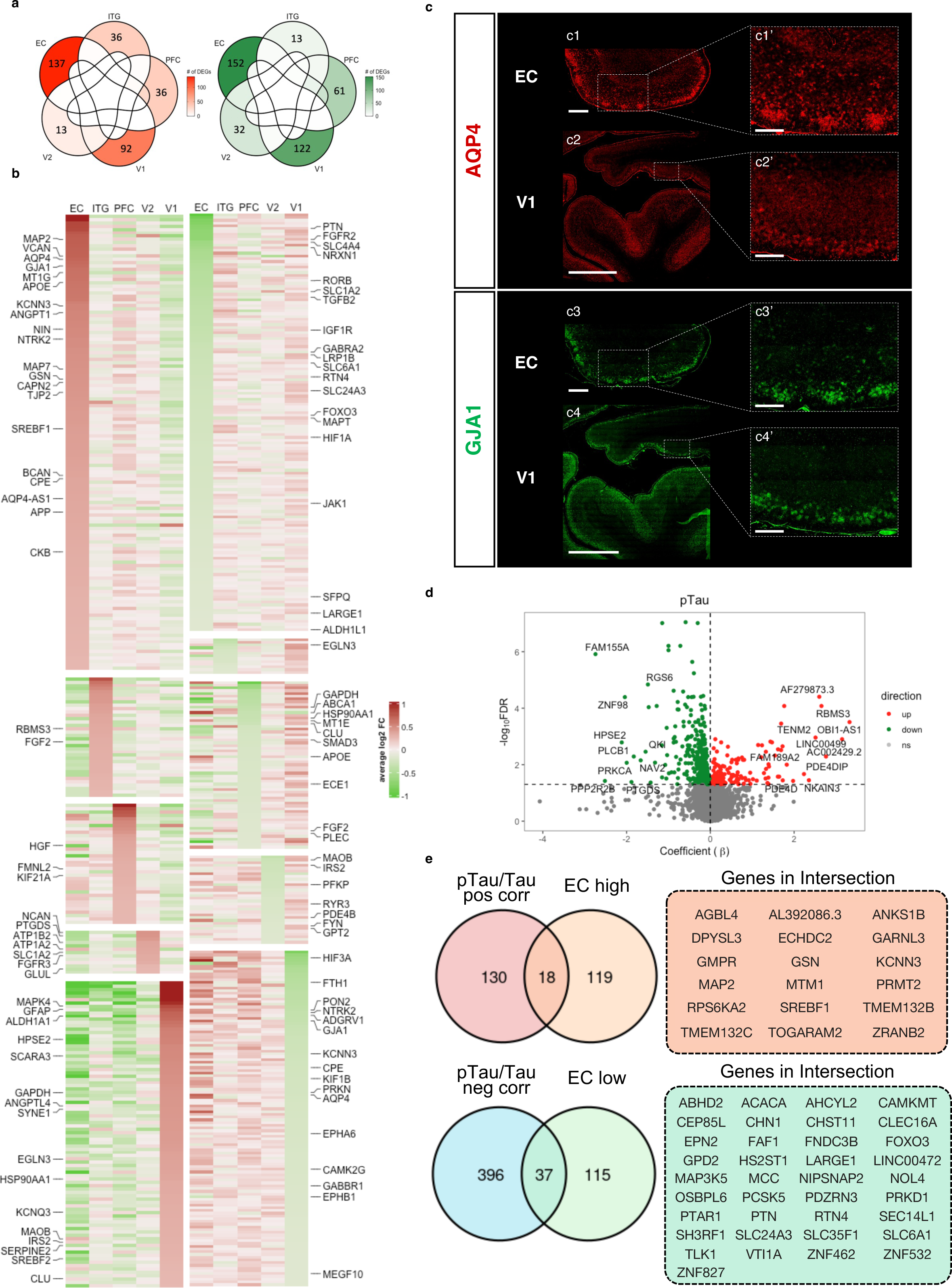
Regional heterogeneity of astrocyte transcriptome in the normal aging brain. **a**, Venn diagrams show the differentially expressed genes (DEGs)—upregulated in red and downregulated in green—in each brain region relative to all other brain regions in n=8 donors at pathology stage 1 (no neuritic plaques and Braak NFT stage 0/I/II). **b**, Heatmap depicts the DEGs in each brain region (upregulated in red and downregulated in green) ranked by average log fold-change. **c**, Fluorescent immunohistochemistry of aquaporin-4 (AQP4) and connexin-43 (GJA1) in EC and V1 demonstrates a higher expression of these two proteins in EC vs. V1 in agreement with the snRNA-seq results. Scale bars: EC, 1 mm; V1, 5 mm; insets, 500 μm. **d**, Volcano plot illustrates the correlation analysis between pTau/Tau and gene expression levels in pathology stage 1 astrocytes. The x-axis represents the correlation coefficient β, while the y-axis indicates the false discovery rate (expressed as -log10FDR). Genes in red are positively correlated and those in green are negatively correlated at an FDR<0.05 (-log10FDR > 1.3), whereas genes in gray were statistically not significant. **e**, Venn diagrams show the number of genes upregulated in EC (EC high) and/or positively correlated with the pTau/Tau ratio (top) and the number of genes downregulated in EC (EC low) and/or genes negatively correlated with pTau/Tau ratio (bottom). Genes in each intersection are listed in the boxes.

These results suggest some degree of region-specificity of astrocyte transcriptome in allocortex vs. primary neocortex vs. association neocortex. To further examine this possibility, we asked whether EC DEGs are driven by the EC microenvironment (i.e., allocortex vs. neocortex) or by the amount of local EC pTau pathology present in Braak I and II donors (Figure 1d and Supplementary Table 1). First, we identified a gene set associated with the local pTau burden by regressing gene expression levels against the pTau/Tau ratio across all five brain regions in these eight donors (148 positively and 433 negatively correlated genes, FDR < 0.05, Supplementary Table 3 and volcano plot in Figure 2d). Then, to disambiguate EC-specific vs. pTau-driven genes, we overlapped these pTau-correlated genes with those enriched in the EC (Venn diagrams in Figure 2e). Only 18 out of the 137 EC-upregulated genes were positively correlated with pTau/Tau ratio (notably including cytoskeletal genes such as *MAP2*, and the master transcription factor regulating cholesterol metabolism *SREBF1*), whereas only 37 out of 152 EC-downregulated genes were negatively correlated with pTau/Tau ratio (including extracellular matrix/glycosaminoglycan metabolism genes such as *LARGE1* and *PTN*). Taken together, these results support the existence of an allocortex-specific gene expression profile in EC astrocytes, which is distinct from a pTau signature.

### Astrocyte transcriptomic changes along the stereotypical spatial progression of AD

Next, we examined whether astrocyte gene expression changes track with the stereotypical spatiotemporal progression of AD neuropathology. Our study design with 32 donors along the normal aging-AD continuum and five sites that are hierarchically involved in classic AD enabled us to answer this question and we hypothesized that the magnitude of astrocyte transcriptomic changes would parallel both the regional vulnerability of neural networks to ADNC (spatial progression) and the temporal accrual of ADNC in a given brain region (temporal progression).

First, to determine whether astrocyte transcriptomic changes follow the neural network predilection of AD progression, we rank-ordered the five brain regions based on their known vulnerability to neurofibrillary degeneration^10, 11^ (i.e., EC>ITG>PFC>V2>V1) and performed a differential expression analysis comparing each node of the network with the next (i.e., EC vs. ITG, ITG vs. PFC, PFC vs. V2, and V2 vs. V1), including all 32 donors, and controlling for within-donor correlation. The resulting 515 DEGs between any two “adjacent” network nodes were then grouped into six spatial gene sets following distinct trajectories along the neural network (Figure 3a, Supplementary Table 4). Two spatial gene sets changed their expression level monotonically along the network, either decreasing from EC to V1 (gene set #1) or increasing from EC to V1 (gene set #2); two spatial gene sets had relatively stable expression levels across all brain regions except for a peak of upregulation (gene set #3) or downregulation (gene set #4) at the PFC; the last two spatial gene sets exhibited relatively constant expression levels in EC, ITG, and PFC, but either decreased (gene set #5) or increased (gene set #6) from the PFC to V2 and V1. Thus, this analysis suggested a significant association between astrocyte gene expression and relevant regions that are hierarchically affected by the AD pathophysiological process.

**Figure 3.**
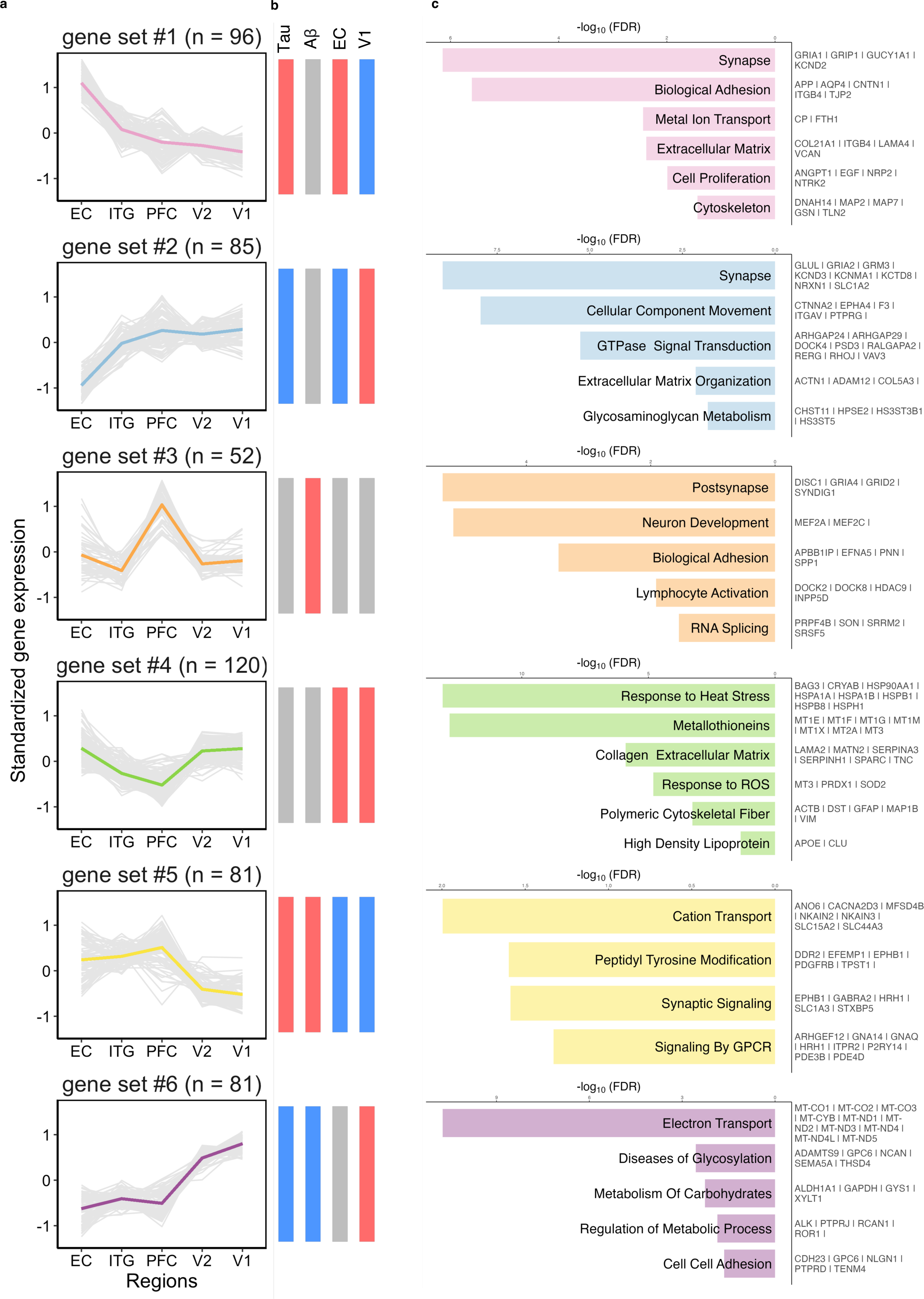
Astrocyte transcriptomic changes along the stereotypical spatial progression of AD. **a**, Spatial trajectory gene sets result from clustering the n=515 DEGs between any two “adjacent” nodes of the AD network from EC to V1. The y-axis is the standardized gene expression with grey lines representing individual genes whereas the colored lines representing the mean trend. **b**, The first two vertical bars represent the association between the average expression of each spatial trajectory gene set in each donor and their pTau/Tau ratio and Aβ plaque burden (red = statistically significant positive correlation; blue = statistically significant positive correlation; gray = non-significant). The last two vertical bars illustrate the results of an overlap test between each spatial trajectory gene set and the region-specific EC and V1 high/low gene sets derived from normal controls in Fig. 2 (red = statistically significant overlap with region-specific upregulated gene set; blue = statistically significant overlap with region-specific downregulated gene set; gray = non-significant). **c**, Functional characterization of each spatial trajectory gene set via pathway analysis; some relevant genes of each pathway are displayed on the right.

To test whether the spatial trajectories of astrocyte gene expression are also related to local A*β* and pTau levels, we correlated the average standardized expression of each of these spatial gene sets for each brain region and donor with the A*β* plaque burden and the pTau/Tau ratio measured in the same brain regions and donors, while controlling for within-donor correlation (Figure 3b, Supplementary Figure 2a). The gene set decreasing linearly from EC to V1 (gene set #1) was positively correlated with the pTau/Tau ratio, whereas the one increasing from EC to V1 (gene set #2) was negatively correlated with the pTau/Tau ratio, suggesting that these two gene sets contain a pTau-associated signature. Gene set #1 contained synaptic (*GRIA1*, *GRIP1*, *KCND2*, *KCNE4*, *KCNN3*, *SYNPO2*, *SYTL4*), cell-cell communication (*APP*, *AQP4*, *CNTN1*, *IL33*, *ITGB4*, *TJP2*), cytoskeleton (*GSN*, *MAP2*, *MAP7*, *MYBPC1*, *TLN2*, *TTN*), extracellular matrix (*ADAMTSL3*, *COL21A1*, *LAMA4*, *SERPINI2*, *VCAN*), and some trophic factors (*ANGPT1*, *EGF*, *NRP2*, *NTRK2*), metal binding (*CP*, *FTH1*), and lipid metabolism (*ABCA1*, *LRAT*) genes, whereas gene set #2 included genes involved in glutamate neurotransmission (*GLUL*, *GRIA2*, *NRXN1*, *SLC1A2*) and extracellular matrix (*ADAM12*, *FREM2*, *MGAT5*, *PTN*) genes (Figure 3c).

By contrast, the gene set specifically upregulated at the PFC (gene set #3) was only positively correlated with the A*β* plaque burden, which was highest at this brain region (Figure 1d), suggesting that this is an A*β*-associated signature. This gene set was enriched in synaptic (*DISC1*, *GRIA4*, *GRID2*, *SYNDIG1*), cell-cell communication genes (*APBB1IP*, *EFNA5*, *EPHA6*, *SPP1*), and cytoskeletal (*SYNM* and *FMN1*) genes that were distinct from those associated with the pTau/Tau ratio, and also contained the GWAS AD risk gene *INPP5D*^14^ and the neuroprotective transcription factors *MEF2A* and *MEF2C*^15^ (Figure 3c). On the other hand, the gene set specifically downregulated at the PFC (gene set #4) was negatively correlated with both A*β* plaque burden and pTau/Tau ratio but the statistical significance was lost after controlling for brain region. This gene set contained important genes involved in cytoskeleton (*ACTB*, *CAP2*, *GFAP*, *MAP1B*, *VIM*), lipid metabolism (*APOE*, *CLU*, *DGKB*), and calcium homeostasis (*CALM2*, *CNN3*, *RTN1*, *S100A1*, *S100B*) as well as many extracellular matrix (*CST3*, *KAZN*, *HS6ST3*, *LAMA2*, *MATN2*, *SDC4*, *SERPINA3*, *SERPINH1*, *SPARC*, *TNC*), proteostasis (*BAG3*, *CRYAB*, *DNAJA1*, *DNAJB1*, *HSP90AA1*, *HSPA1A*, *HSPA1B*, *HSPB1*, *HSPB8*, *HSPD1*, *HSPE1*, *HSPH1*, *UBB*, *UBC*), and antioxidant defense genes (*PRDX1*, *SOD2*, and the metallothioneins *MT1E*, *MT1F*, *MT1G*, *MT1M*, *MT1X*, *MT2A*, *MT1M*, *MT2A*, *MT3*) (Figure 3c).

Lastly, spatial gene set #5 was positively correlated with both pTau/Tau ratio and A*β* plaque burden, whereas gene set #6 was negatively correlated with both measures, likely representing ADNC-associated pan-reactive upregulated and downregulated signatures, respectively. The pan-reactive genes positively correlated with pTau and A*β* levels included many genes involved in G-protein-coupled receptor (GPCRs) signaling (*ARHGEF12*, *GNA14*, *GNAQ*, *PDE3B*, *PDE4D*, *PDE4DIP*, *PDE5A*, *PRKG1*) and intracellular transport (*CPQ*, *DNAH7*, *DNM3*, *RANBP3L*, *SLC1A3*, *SLC15A2*, *SLC44A3*, *SLCO1C1*), suggesting an activation of second messenger-mediated signaling cascades and intracellular trafficking of solutes and vesicles (Figure 3c). Remarkably, the pan-reactive genes negatively correlated with pTau and A*β* levels were predominantly glucose metabolism (*ALDH1A1*, *DPP6*, *GAPDH*, *GYS1*, *LDHB*) and mitochondrial electron transport chain (*MT-ATP6*, *MT-CO1* to *MT-CO3*, *MT-CYB*, *MT-ND1* to *MT-ND5*, *MT-ND4L*) genes, suggesting a failure of energy metabolism in reactive astrocytes associated with chronic exposure to both pTau and A*β* (Figure 3c).

We then asked whether this association could be driven by the regional differences observed in normal control donors or whether it could be explained by the accumulation of A*β* and/or pTau in those brain regions. To test the first possibility, we compared the EC-specific and V1-specific signatures obtained from normal controls (Figure 2) and these spatial trajectory gene sets derived from the entire sample and, indeed, observed a significant overlap (Supplementary Figure 2b). Specifically, the gene set with monotonic decrease from EC to V1 (gene set #1) was significantly enriched in EC-high and V1-low genes, whereas the gene set with monotonic increase from EC to V1 (gene set #2) was significantly enriched in EC-low and V1-high genes; the gene set specifically upregulated in PFC (gene set #3) was not enriched in either EC or V1 signatures, whereas the gene set specifically downregulated in PFC was enriched in EC-high and V1-high genes; and the gene set with relatively stable expression in EC through PFC and lower expression in V2 and V1 (gene set #5) was enriched in EC-low and V1-low genes; and the gene set with stable levels in EC through PFC but increase in V2 and V1 (gene set #6) was expectedly enriched in V1-high genes (Figure 3b). These data suggest that some of the regional variations in astrocyte transcriptome along the AD neural network are already present in the normal aging brain, possibly driven by exposure of astrocytes to microenvironmental factors particular to each brain region.

Taken together, these results indicate that the astrocyte gene expression profile parallels the typical spatial progression of AD pathology along neural networks and suggest that these astrocyte transcriptomic changes are partly associated with the local levels of pTau and/or A*β* plaques, and partly explained by region-specific microenvironmental factors.

### Astrocyte transcriptomic changes along the temporal progression of AD

Once we established the spatial progression of astrocyte transcriptomic changes in the normal aging-AD continuum, we sought to determine whether astrocyte transcriptomic changes also parallel the temporal accrual of ADNC within a given brain region. To this end, we conducted a differential expression analysis comparing each of the aforementioned four pathology stages with the next, while controlling for brain region and within-donor correlation. We decided to separate Braak V and VI donors to investigate possible end-stage changes in astrocyte transcriptome and because, by definition, the primary visual cortex (V1) only contains NFTs in Braak VI donors but is spared by NFTs in Braak V donors^10, 11^. The resulting 806 DEGs between any two “adjacent” stages were, thus, considered temporally associated with AD pathophysiology. These 806 DEGs were grouped into six different temporal gene sets with distinct temporal trajectories and variable strength of association with the local A*β* and pTau levels (Figure 4, Supplementary Table 5).

**Figure 4.**
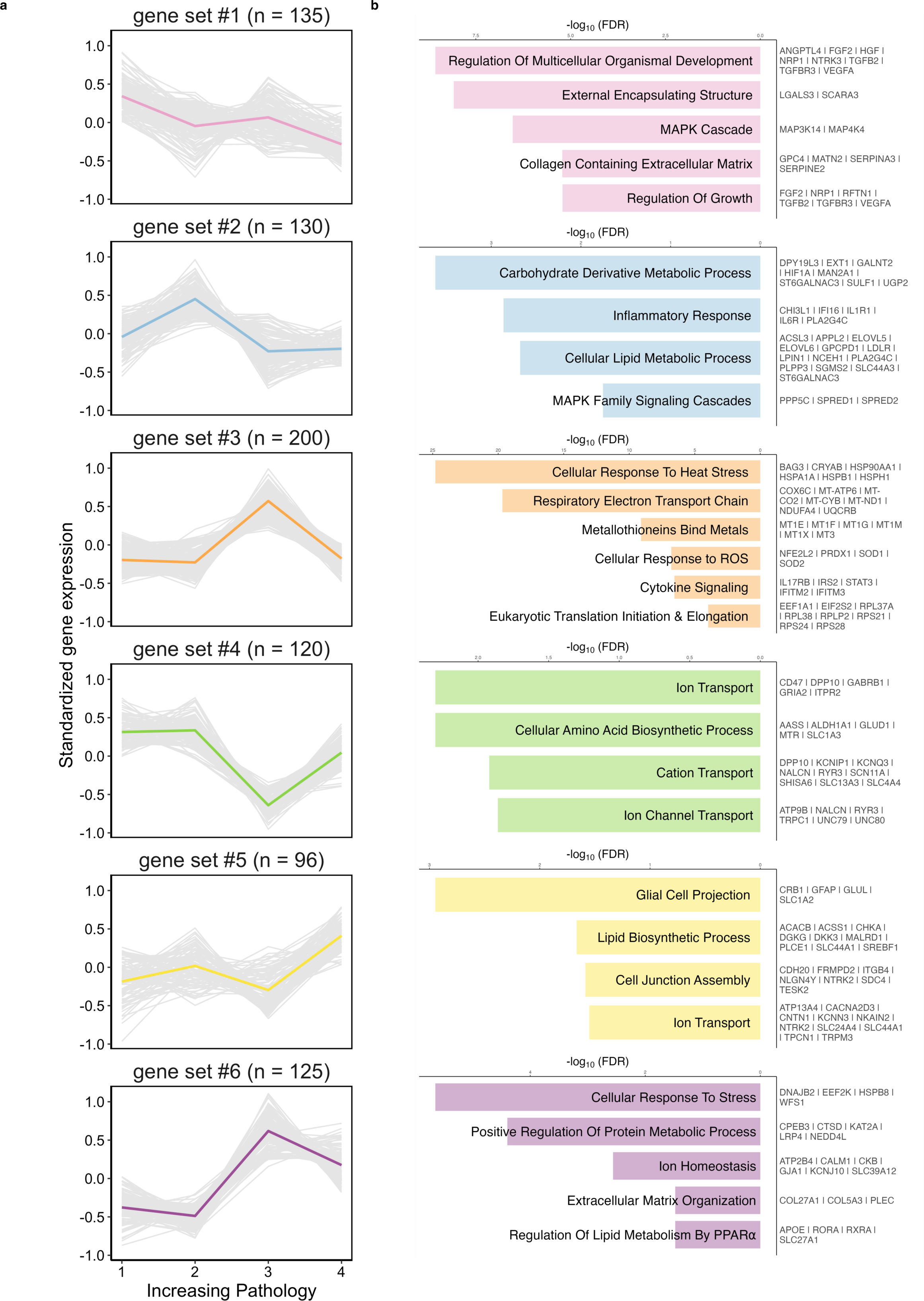
Astrocyte transcriptomic changes along the temporal progression of AD. **a**, Temporal trajectory gene sets result from clustering the n=806 DEGs between any two “adjacent” pathology stages from early to end-stage. The y-axis is the standardized gene expression with grey lines representing individual genes whereas the colored lines represent the mean trend. **b**, Functional characterization of each temporal trajectory gene set via pathways analysis; some relevant genes of each pathway are displayed on the right.

One set (gene set #1) had the highest expression at early stage (no NPs, Braak NFT stage 0/I/II), then decreased and remained relatively stable along all the other stages, suggesting that it corresponds with a homeostatic signature of astrocytes. This gene set included many trophic factors and their receptors (*ANGPTL4*, *APC*, *EGLN3*, *FGF2*, *GFRA1*, *HGF*, *IGF1R*, *NRP1*, *NTRK3*, *TGFB2*, *TGFBR3*, *VEGFA*), extracellular matrix (*GPC4*, *MATN2*, *SERPINA3*, *SERPINE2*, *SPOCK1*), cytoskeleton (*ACTN1*, *FIGN*, *MACF1*, *MAPRE2*, *MAST4*, *MYH9*), and some metal-binding (*CP*, *FTL*, *MT2A*), anti-oxidant (*MGST1*, *MSRA*) but also pro-oxidant enzymes (*MAOB*), neuroinflammation (*MAP3K14*, *MAP4K4*, *OSMR*, *SOCS3*, *TAB2*), calcium homeostasis (*CALN1*, *CNN3*, *S100A1*), and phagocytosis (*LGALS3*, *SCARA3*) genes.

Another (gene set #2) peaked at the intermediate ADNC stage (sparse/moderate neuritic plaques and Braak NFT stage III) and then returned to baseline at late and end-stages, suggesting a transient activation of this gene set in response to the initial accumulation of ADNC. This early gene program included trophic and survival factors (*FGF1*, *FGF14*, *HIF1A*, *NRP2*), extracellular matrix (*GPC6*, *KAZN*, *LAMA4*, *TNC*), cytoskeleton (*ABI1*, *CLIP1*, *MYO1E*, *VCL*), and neuroinflammation (*CHI3L1*, *IFI16*, *IL1R1*, *IL6R*, *RCAN1*) genes different from those predominant in the early ADNC stage, plus lipid metabolism (*ACSL3*, *ELOVL5*, *ELOVL6*, *LDLR*, *LEPR*, *LPIN1*, *NCEH1*, *OSBPL6*, *PLA2G4C*, *PLCL1*, *PLPP3*, *SGMS2*), glycosaminoglycan metabolism (*EXT1*, *GALNT2*, *MAN2A1*, *ST6GALNAC3*, *SULF1*, *UGP2*), synaptic (*GRIK2*, *PCLO*, *STXBP6*, *SYTL2*), and GPCRs-mediated signaling (*GPR158*, *KALRN*, *PDE4B*, *PDE4D*, *RASGEF1B*, *RGS6*, *TRIO*) genes.

Two other temporal gene sets either peaked (gene set #3) or dropped (gene set #4) at the late stage (moderate/frequent neuritic plaques, Braak NFT stage V) but surprisingly returned to (near) baseline at end-stage (moderate/frequent neuritic plaques, Braak NFT stage VI), suggesting a vigorous response of astrocytes to nearby A*β* plaques and NFTs that they ultimately cannot sustain and turn off as this exposure becomes chronic. The upregulated late gene program was comprised of genes involved in proteostasis/response to heat stress (*AHSA1*, *BAG3*, *CCT4*, *CRYAB*, *DNAJA1*, *DNAJB1*, *DNAJB6*, *HSP90AA1*, *HSP90AB1*, *HSPA1A*, *HSPA1B*, *HSPA4*, *HSPA4L*, *HSPA8*, *HSPA9*, *HSPB1*, *HSPD1*, *HSPE1*, *HSPH1*, *MIB1*, *OTUD7B*, *SQSTM1*, *ST13*, *UBB*, *UBC*, *UBE2B*, *UBE2E2*), protein translation (*EEF1A1*, *EIF1*, *EIF2S2*, *RPL37A*, *RPL38*, *RPLP2*, *RPS21*, *RPS24*, *RPS28*), energy metabolism (*ENO1*, *GAPDH*, *LDHB*, *PGK1*), mitochondrial electron transport chain (*ATP5F1E*, *ATP5MD*, *ATP5ME*, *ATP5PF*, *COX6A1*, *COX6C*, *COX7C*, *MT-ATP6*, *MT-CO2*, *MT-CO3*, *MT-CYB, MT-ND1*, *MT-ND2*, *MT-ND3*, *MT-ND4*, *NDUFA4*, *NDUFB2*, *NDUFB4*, *NDUFC1*, *UQCRB*), antioxidant defense (*NFE2L2*, *NXN*, *PRDX1*, *SOD1*, *SOD2*, and the metallothionein genes [*MT1E*, *MT1F*, *MT1G*, *MT1M*, *MT1X*, *MT3*]), neuroinflammation (*IFITM2*, *IFITM3*, *IL17RB*, *IRS2*, *NFAT5*, *PTGES3*, *STAT3*), cell-cell communication (*CD59*, *CDH23*, *CHL1*, *CNTNAP2*, *CNTNAP3*, *CNTNAP3B*, *FGFR1*, *RGMA*), and some genes involved in lipid metabolism (*ABHD3*, *CLU*, *OAZ1*, *OSBPL1A*), cytoskeleton (*CLIP2*, *MAP2*, *SYNM*, *VIM*), extracellular matrix (*MMP16*, *PLOD2*, *PLOD3*, *SERPINH1*, *SPP1*, *ST6GALNAC6*), synaptic function (*CAMK2D*, *CAMK2N1*, *FOS*, *GAB1*, *NRXN3*, *RIMS1*, *SYT1*), and intracellular transport/trafficking (*ATP2C1*, *CPE*, *DYNLRB1*, *SLC7A5*, *SLC7A11*, *SLC9B2*, *SLC20A2*, *SNX3*). The downregulated late gene program included genes involved in glutamate neurotransmission (*GLUD1*, *GRIA2*, *SLC1A3*), extracellular matrix (*COL28A1*, *CSGALNACT1*, *EPM2A*, *GPC5*), and intracellular transport (*GOLGA8B*, *SLC4A4*, *SLC13A3*, *TVP23C*). Interestingly, many of these late-stage genes overlap with the A*β*/pTau-unrelated genes (gene set #4) of the spatial progression analysis. This lack of correlation with A*β* and pTau is likely explained by the “normalization” of their expression levels in end-stage disease (Braak VI) despite further accumulation of A*β* and especially pTau in all brain regions of Braak VI donors.

Moreover, we identified a smaller set (gene set #5) whose expression levels remained relatively stable throughout early, intermediate, and late stages, and only increased at end-stage (moderate/frequent neuritic plaques, Braak NFT stage VI); this gene set contained glutamate metabolism (*GLUL*, *SLC1A2*), extracellular matrix (*ADAMTSL3*, *COL21A1*, *HPSE2*, *SDC4*, *VCAN*), lipid metabolism (*ACACB*, *ACSS1*, *DGKG*, *OSBPL11*, *PHYHD1*, *PHYHIPL*, *PLCE1*, *PPARGC1A*, *SREBF1*), trophic factors (*FGFR3*, *NTRK2*), cell-cell communication (*CDH20*, *CLEC16A*, *CNTN1*, *ITGB4*, *NLGN4Y*), and intracellular transport (*DYNC2H1*, *SCLT1*, *SLC14A1*, *SLC18B1*, *SLC24A4*, *SLC44A1*) genes. Lastly, gene set #6 had low expression at early and intermediate stages, peaked at late stage, and decreased at end-stage without returning to baseline; relevant genes pertained to proteostasis (*CTSD*, *DNAJB2*, *HSPB8*, *NEDD4L*), energy metabolism (*ALDH2*, *CKB*, *PFKP*), extracellular matrix (*B4GALNT4*, *COL5A3*, *COL27A1*, *CST3*, *FLRT2*, *PLEC*, *PLXNB1*), intracellular transport and trafficking (*ATP2B4*, *SLC27A1*, *SLC38A2*, *SLC39A11*, *SLC39A12*, *TRAK1*), cell-cell communication (*GJA1*), lipid metabolism (*APOE*, *LRP4*), and nuclear receptors (*RORA*, *RXRA*).

Taken together, these results indicate that both brain region and Braak NFT stage should be carefully considered when comparing the astrocyte transcriptome of AD vs. normal control brains because astrocyte transcriptomic responses to ADNC might be attenuated in end-stage disease (Braak NFT stage VI) in severely and chronically affected brain areas.

### Astrocyte clustering analysis reveals diverse transcriptomic programs across brain regions and donors

While two broad categories of astrocytes, homeostatic and reactive, have been traditionally postulated, the above results suggested a more complex picture of astrocyte reactivity^2^. Thus, we investigated the existence of distinct subpopulations or states of astrocytes based on their gene expression programs. After removing donor-specific subclusters and subclusters containing multiplets with other cell types, clustering of astrocyte nuclei based on their transcriptome at 0.3 resolution rendered six astrocyte subclusters in ITG, PFC, and V2, and eight in EC and V1. To correlate these subclusters across brain regions, we performed a Spearman’s rank correlation with the rank values of the genes defining each subcluster in each brain region. This analysis identified a total of 10 astrocyte subclusters across the five brain regions (Figure 5a). Inspection of both the top DEGs defining these 10 subclusters (i.e., marker genes for these subclusters) and the enriched pathways led us to define two homeostatic subclusters (astH1 and astH2) and two reactive subclusters (astR1 and astR2), with six additional “intermediate” subclusters (astIM1 to astIM6).

**Figure 5.**
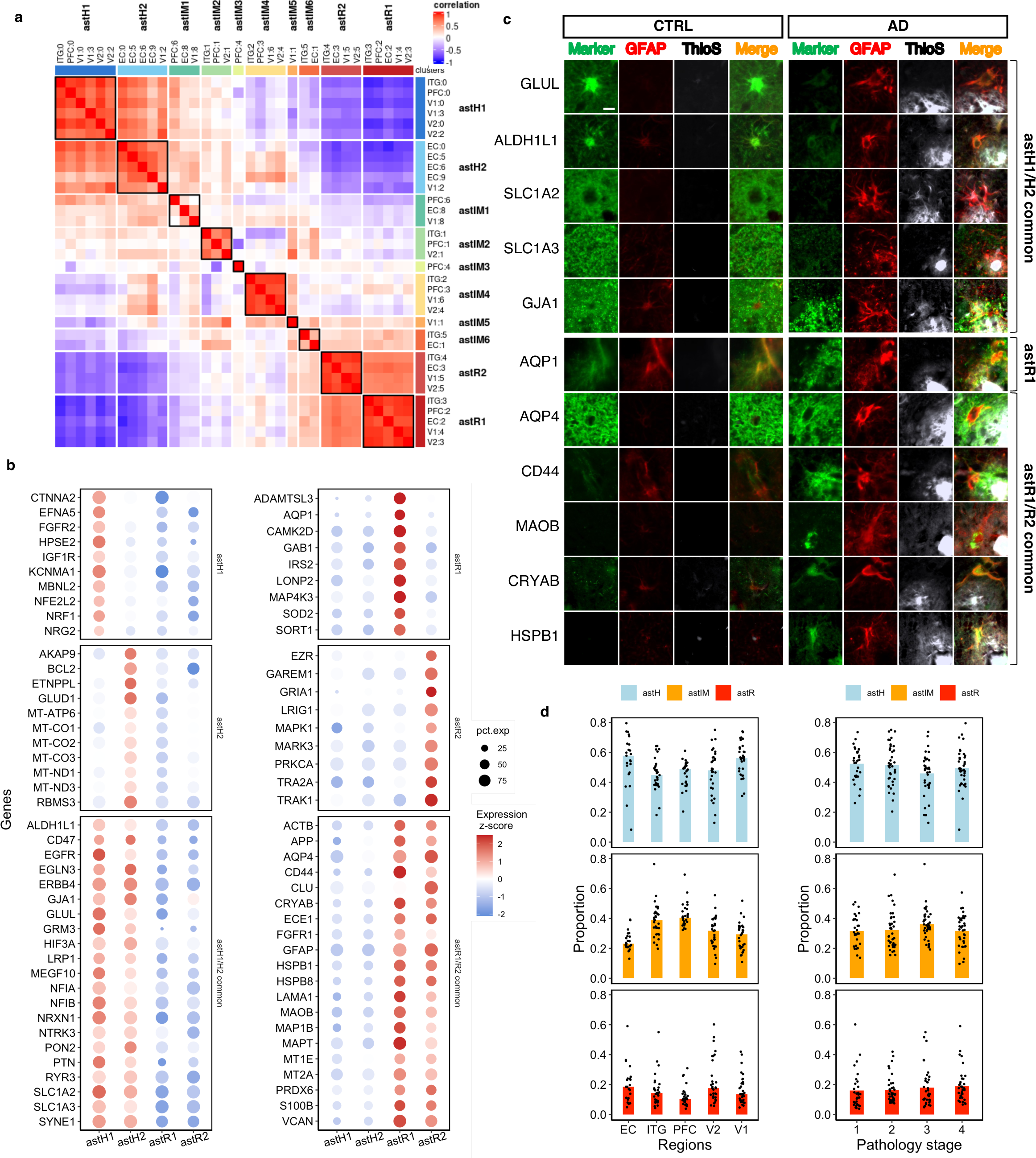
Clustering reveals homeostatic, reactive, and intermediate astrocytes. **a**, Heatmap of spectral clustering mapping astrocytes subclusters across brain regions based on the strength of the correlation of their gene expression results in 10 main astrocyte clusters. **b**, Bubble plots illustrate the expression z-scores of selected marker genes defining the two homeostatic clusters (astH1 and astH2) and the two reactive clusters (astR1 and astR2). **c**, Double fluorescent immunohistochemistry for selected markers with Thioflavin-S (ThioS) counterstaning in formalin-fixed paraffin-embedded sections from the temporal association cortex of control (CTRL) and AD donors show reduced expression of some homeostatic markers and increased expression of some reactive markers in GFAP+ astrocytes surrounding some ThioS+ Aβ plaques. Note: photomicrographs from CTRL and AD donors were taken with similar exposure time and display settings for appropriate comparison. Scale bar: 10 μm. **d**, Proportions of astrocytes clusters across brain regions and pathology stages (astH = astH1 + astH2, astIM = astIM1 + astIM2 + astIM3 + astIM4 + astIM5, astR= astR1 + astR2).

We found that astH1 and astH2 were highly correlated (*ρ* > 0.8), with common genes including *ALDH1L1*, genes encoding growth factor receptors (*EGFR*, *NTRK3*), and genes involved in cell-cell adhesion (*ERBB4*, *NRXN1*), phagocytosis (*CD47*, *MEGF10*), and glutamatergic neurotransmission (*GRM3*, *GLUL*, *SLC1A2*, *SLC1A3*) (Figure 5b). Marker genes enriched in astH1 included growth factor receptors (*FGFR2*, *IGF1R*, *NRF1*) and the antioxidant transcription factor *NFE2L2*, whereas marker genes of astH2 included many mitochondrial electron transport chain genes (*MT-ATP6*, *MT-CO1*, *MT-CO2*, *MT-CO3*, *MT-ND1*, *MT-ND3* and the anti-apoptotic gene *BCL2* (Figure 5b). Reactive astrocyte clusters astR1 and astR2 were also highly correlated with each other (*ρ* > 0.8) and both expressed *APP*, *AQP4*, heat shock proteins (*CRYAB*, *HSPB1*, *HSPB8*), metallothioneins (*MT1E*, *MT2A*), extracellular matrix (*CD44*, *LAMA1*, *VCAN*), cytoskeletal proteins (*ACTB*, *GFAP*, *MAP1B*, *MAP2*, *MAPT*), and *MAOB* (Figure 5b). Marker genes enriched in astR1 included *AQP1*, *SOD2*, and *SORT1*, whereas *CLU*, *GRIA1*, and *MAPK1* among others defined astR2 (Figure 5b). Fluorescent immunohistochemistry shows that many of these astR1/R2 markers are highly expressed by GFAP+ve reactive astrocytes near Thioflavin-S+ve dense-core A*β* plaques (Figure 5c and Supplementary Figures 3 and 4). Fluorescent *in situ* hybridization via RNAScope confirmed *MAPT* expression in astrocytes (Supplementary Figure 5).

Pathway enrichment analysis against the Reactome database revealed that, compared to all other astrocyte subclusters, astH1 upregulate neurotransmission, glycosaminoglycan metabolism, and cell-cell interaction via tight junctions, and both astH1 and astH2 downregulate the HSF1-mediated heat shock response, whereas no Reactome pathway was statistically significantly enriched in upregulated astH2 genes. By contrast, astR1 upregulate metallothioneins, aquaporin-mediated transport, and carbohydrate metabolism and down-regulate neurotransmission; astR2 upregulate phagocytosis uptake, NLRP3 inflammasomes, and lipid metabolism, and both upregulate HSF1-mediated heat shock protein response, extracellular matrix, and toll-like receptor cascades, and downregulate cell-cell interaction via tight junctions and ephrin signaling (Supplementary Table 6).

While at least three major astrocyte subclusters (homeostatic, reactive, and intermediate) could be differentiated based on their transcriptome, surprisingly, the proportion of nuclei classified within these three broad categories did not differ across pathology stages (Figure 5d), with roughly 10-25% of astrocyte nuclei classified as astR1 or astR2, 45-60% as astH1 or astH2, and 25-45% as astIM1 to astIM6. Moreover, although these three broad subclusters also had similar proportions across brain regions, some regional differences were evident when comparing astH1 vs. astH2 and astR1 vs. astR2: astH2 were only present in EC and V1, whereas astR2 were absent in PFC. Taken together, these data indicate that there are very distinct transcriptomic subclusters of astrocytes with unexpectedly similar proportions throughout the cortex in both normal aging and AD brains.

### Intermediate astrocytes are transitional states between homeostatic and reactive astrocytes

Lastly, we asked whether these homeostatic and reactive astrocyte subclusters represent stable astrocyte subpopulations or more transient states comprising a dynamic astrocyte gene expression continuum. A closer inspection of the rank correlations across astrocyte subclusters shown in Figure 5a indicated that the intermediate subclusters astIM1 and astIM2 are more similar to the homeostatic subclusters (astH1 and astH2), whereas the intermediate subclusters astIM5 and astIM6 are more similar to the reactive subclusters (astR1 and astR2). In addition, the pathway enrichment analysis of the intermediate astrocyte subclusters (astIM1 through astIM6) revealed a gradient of upregulated and downregulated enriched pathways from homeostatic to reactive astrocytes (Supplemental Table 6). Moreover, a pseudotime trajectory analysis in a subsample of astrocytic nuclei randomly selected from all donors and all brain regions identified six distinct possible gene expression trajectories between astH1 and astR1 subclusters, which tracked through astIM subclusters (Figure 6a-c, Supplementary Table 7). Representative genes following each trend are shown in Figure 6d. Taken together, these findings support the idea that homeostatic and reactive astrocytes are dynamic transcriptional states of the same astrocytes rather than different subpopulations of astrocytes, and that intermediate astrocytes represent transitional states between homeostatic and reactive.

**Figure 6.**
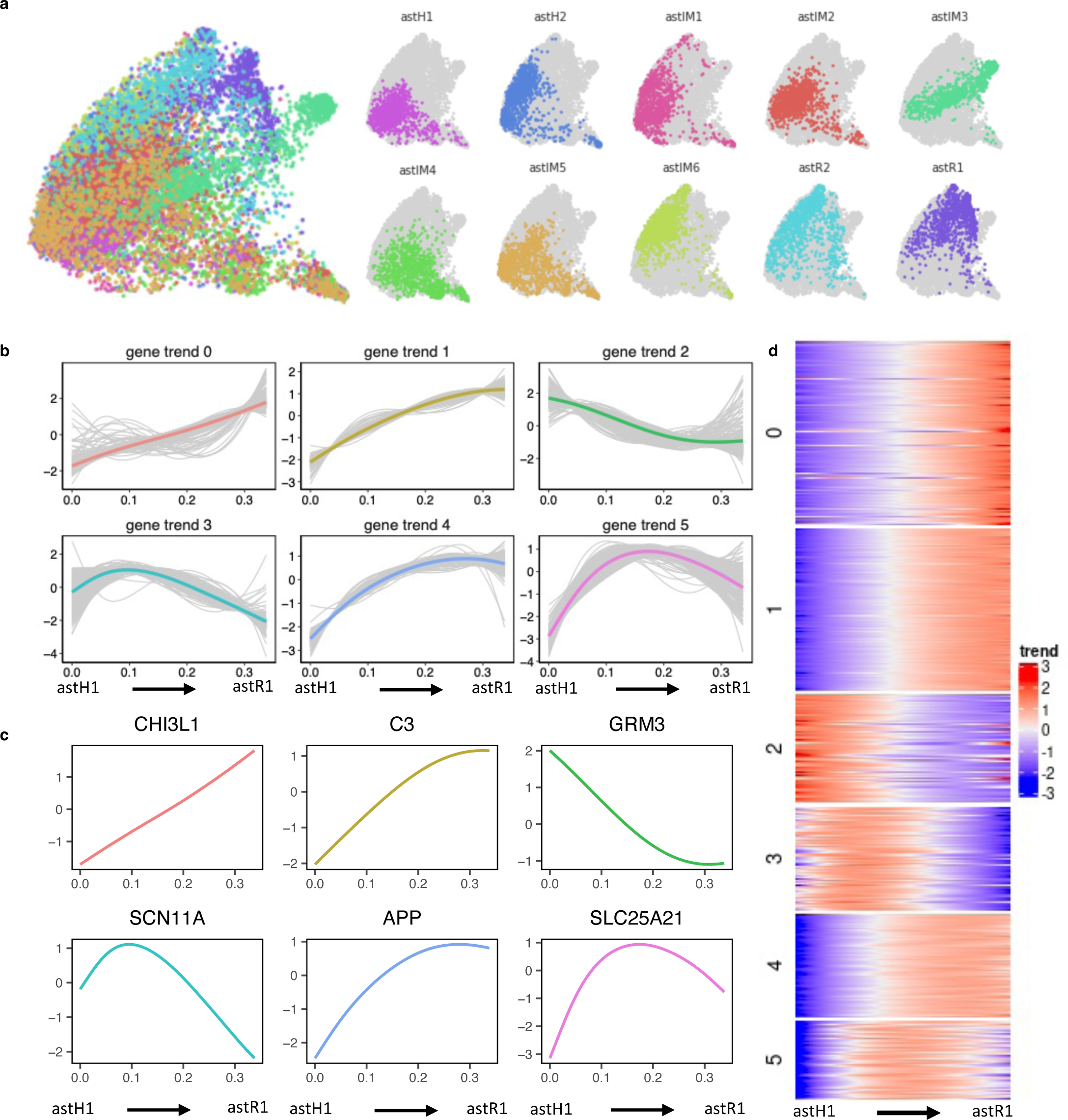
Pseudotime analysis of homeostatic, reactive, and intermediate astrocytes reveals that these are astrocyte states rather than separate subpopulations. **a**, UMAP plot of a subsample of astrocyte nuclei from all 32 donors included in the pseudotime analysis with the colors representing each of the 10 astrocyte subclusters resulting from the clustering in Fig. 5. Note that astH and astR nuclei are most distant, with astIM nuclei mapping in between. **b**, Distinct gene trends resulting from the pseudotime analysis between astH1 and astR1. **c**, Representative genes from each of these pseudotime trends between astH1 and astR1. **d**, Heatmap illustrates the gene sets following each of these six pseudotime trends between astH1 and astR1.

## DISCUSSION

We generated the largest snRNA-seq dataset to date of human astrocytes across five brain regions and encompassing the normal aging-AD continuum. The analysis of this large dataset provided important clues on the regional heterogeneity of astrocyte transcriptome in the normal aging brain and the transcriptomic programs that astrocytes activate and deactivate in the AD brain in response to the stereotypical spatiotemporal progression of ADNC.

First, we observed significant regional differences in astrocyte transcriptome in the brains from normal controls, with EC and V1 region showing the highest numbers of upregulated and downregulated DEGs. The EC, specifically layer II, is the first region affected by NFTs and neurodegeneration in most AD cases^10, 16^. Here we identified an EC-specific astrocyte signature, which included changes in genes relevant to AD pathophysiology such as an upregulation of *APP* and *APOE* and a downregulation of *SLC1A2* (encoding the glutamate transporter EAAT2, a.k.a. GLT-1) and *MAPT* (encoding the microtubule associated protein tau). Noteworthy, an increase in *SLC1A2* expression in EC astrocytes has been associated with resilience to AD pathology^17^, and downregulation of *SLC1A2* is predicted to increase the excitability of glutamatergic neurons^18^, suggesting a potential role of astrocytic genes in EC vulnerability signatures. Also of note, we unexpectedly found robust expression of *RORB* in astrocytes, although *RORB* had previously been identified by snRNA-seq as a marker gene for excitatory EC neurons vulnerable to NFTs and neurodegeneration in AD^5^. By contrast, our study identified *RORB* as one of the downregulated genes in EC astrocytes, arguing for future investigation into its cell type-specific functions. The virtual absence of A*β* deposits across brain regions in these control donors enabled us to isolate the effects of pTau pathology on the astrocyte transcriptome and to confirm that the EC signature cannot be explained just by the NFTs already present in the EC in donors with Braak NFT stages I and II. On the other hand, the V1 signature in control donors could be, at least partly, mediated by the astrocytes from the *stria* of Gennari or Vicq-d’Azyr, a band of white matter that lies between layers IVb and IVc of the calcarine cortex and defines the striate cortex of BA17^19^, since white matter fibrous astrocytes are markedly different from protoplasmic cortical astrocytes^20, 21^. Thus, the substantial regional heterogeneity found in the normal aging brain warrants careful interpretation of region-specific studies comparing AD vs. control donors.

Second, we leveraged of our multi-region sampling of many donors with various levels of ADNC to investigate associations between astrocyte gene expression levels and the spatiotemporal progression typical of AD. When we rank-ordered the five brain regions along a spatial axis following the typical hierarchical progression of AD and compared the astrocyte transcriptome from all donors across any two “adjacent” nodes of the AD network, we were able to identify several gene sets following distinct spatial trajectories. We found that these various spatial trajectories of astrocyte gene expression could only be partly explained by the regional differences detected in the normal controls with virtually no A*β* plaques and pTau pathology mostly restricted to the EC. On the contrary, except for one gene set, all other spatial trajectories were also associated with the local levels of either A*β*, pTau, or both, enabling us to investigate A*β*- and pTau-associated gene signatures. A*β* and pTau levels positively correlated with different synaptic, cell-cell communication, and cytoskeletal genes. Of note are genes involved in the astrocytic component of tripartite synapses that are dysregulated in parallel to the stereotypical march of pTau pathology across cortical association areas. Furthermore, the negative correlation between the *SLC1A2-*containing synapse-enriched spatial gene set and pTau levels is consistent with the reported association between a PET-based tau-spreading network and a gradient of expression of glutamatergic synaptic genes, singularly *SLC1A2*^22^. Thus, these results could be indicating that astrocytes, as part of the tripartite excitatory synapses, play a key role in synaptic dysfunction and the trans-synaptic pTau propagation between excitatory neurons along vulnerable AD neural networks. On the other hand, genes correlated with both A*β* and pTau levels (a.k.a. pan-reactive) consisted of an upregulation of GPCRs-mediated signaling and intracellular trafficking, and a downregulation of glucose metabolism and mitochondrial electron transport, suggesting a synergistic effect of A*β* and pTau on the astrocyte transcriptome^23^. The latter result also suggests that astrocytes undergo an energy failure as a result of the chronic exposure to both A*β* and pTau. Notably, prior bulk transcriptomic analyses have highlighted energy metabolism deficits as a feature of AD and other neurodegenerative diseases^23^ but have either attributed this finding to the severe neuronal dysfunction and loss or imputed it to astrocytes only through indirect computational methods^24^. Our data seem to confirm that astrocytes, and not only neurons, suffer mitochondrial dysfunction and energy deficits that contribute to these observed bulk RNA changes. It should be noted that some of these genes are mitochondrial—rather than nuclear— encoded genes, potentially indicating non-nuclear ambient RNA contamination. However, ambient RNA contamination is mainly of neuronal origin and substantially reduced by depletion of neuronal nuclei^25^ and the expression of these mitochondria-encoded genes was co-regulated with nuclear-encoded mitochondrial genes, favoring the conclusion that these findings represent true biological changes. Intriguingly, astrocytes have been reported to contribute to the [^18^F]FDG-PET signal via the glutamate transporter GLT-1, encoded by *SLC1A2*^26^ (whose expression is negatively associated with pTau). Also interestingly, higher plasma GFAP levels have been associated with higher [^18^F]FDG-PET signal in early AD^27^. Thus, reactive astrocytes may initially increase their metabolic rate but finally fail to meet their energy needs and contribute to the typical bilateral temporo-parietal hypometabolism observed in AD. Interestingly, another sizable gene set (gene set #4, n =120), characterized by its lack of correlation with either A*β* or pTau levels and highest expression levels in the EC and V1, was composed of genes related to the cytoskeleton (including *GFAP*), lipid metabolism (including *APOE* and *CLU*), calcium homeostasis, extracellular matrix, heat shock proteins, and antioxidant defense (e.g., metallothioneins), suggesting a non-linear heterogenous response to ADNC in the different cortical regions.

We then examined the astrocyte transcriptomic changes along the temporal axis by analyzing the effect of increasing ADNC severity, which matched well with increasing local levels of A*β* and pTau in each brain region. Assuming that autopsy findings of ADNC severity can be temporally ordered, which is well supported by the PET imaging biomarker literature, this analytical approach revealed what appears to be successive waves of astrocyte transcriptomic programs in intermediate vs. late vs. end-stage disease. At the intermediate ADNC stage, astrocytes induced trophic and survival factors (significantly the transcription factor hypoxia-inducible factor-1*α* gene *HIF1A*, which has been shown to be a major driver of energy metabolism changes in AD reactive microglia^28^), extracellular matrix (e.g., *GPC6* and *TNC*), cytoskeleton, and neuroinflammation (notably the gene *CHI3L1* encoding for the fluid biomarker of reactive astrogliosis YKL-40 ^29^ and the endogenous inhibitor of calcineurin *RCAN1*) genes different from those predominant in the earliest ADNC stage. Additionally, lipid metabolism (notably the APOE receptor *LDLR*^30^ and the fatty acid elongases *ELOVL5* and *ELOVL6*, which could result in the generation of purportedly toxic saturated long-chain fatty acids^31^), glycosaminoglycan metabolism, synaptic, and GPCRs-mediated signaling genes were also upregulated in the intermediate stage. We separated Braak V and VI donors to test the hypothesis that gene expression by reactive astrocytes may be impacted by chronic exposure to A*β* and pTau, represented by this disease end-stage. Indeed, the distinguishing feature between Braak NFT stages V and VI is not just the presence of NFTs in the primary visual (V1), motor, and sensory cortical areas, but more severe neurofibrillary degeneration^11^ and atrophy throughout the cortical mantle^32^. We observed that at the late stage (Braak V), the astrocyte transcriptomic response was dominated by an upregulation of genes related to protein translation (i.e., elongation factors and ribosomal subunits), proteostasis (i.e., many small chaperones and heat shock proteins), energy metabolism (e.g., *GAPDH* and *LDHB*), mitochondria electron transport chain, neuroinflammation (notably interferon response genes and *STAT3*), anti-oxidant defense (e.g., *SOD1*, *SOD2*, and many metallothionenin genes), and the cytoskeleton (e.g., *SYNM* and *VIM*), suggesting a full-blown, “all-hands-on-deck” adaptive response to nearby pervasive and conspicuous ADNC. Surprisingly, we observed that many of the genes upregulated during late stages (Braak V) decreased or even returned to “baseline” during end-stage disease (Braak VI), including those involved in proteostasis (e.g., heat shock response), energy metabolism and mitochondrial oxidative phosphorylation, lipid metabolism (significantly *APOE* and *CLU*), cytoskeleton, and extracellular matrix, suggesting an exhaustion of astrocyte response. Many of the genes blunted in end-stage overlapped with the A*β*/pTau-uncorrelated genes (gene set #4) found in the spatial progression analysis, thereby explaining their lack of correlation with A*β* and pTau levels. We note that this “burnt-out” phase of astrocyte response in Braak VI donors has not been previously described and may have confounded previous studies that compared only control and severe AD donors or grouped late- and end-stage donors (Braak V/VI).

Taken together, these findings reinforce the idea that AD-associated reactive astrogliosis tracks with the spatiotemporal progression of AD, highlighting a severe end-stage dysfunctional or maladaptive response of astrocytes likely caused by their chronic exposure to A*β*, pTau, and ongoing neurodegeneration^24^.

Third, we identified distinct subclusters of homeostatic and reactive astrocytes based on their gene expression profiles and investigated possible subcluster shifts along the spatiotemporal progression of AD. While we did find some regional specificity of particular subclusters of homeostatic and reactive astrocytes, surprisingly, we found no major differences in the proportion of these subclusters along the temporal axis indicated by ADNC severity stages. This is in agreement with some prior snRNA-seq studies^5, 7^ (whereas others did find one or more subclusters of AD-associated astrocytes^4, 6, 33^) and could be explained by limitations inherent to the clustering method of dimensionality reduction^34^, technical difficulty in isolating nuclei from GFAP-laden astrocytes causing selection bias against the most reactive astrocytes, differences between nuclear vs. cytosolic RNA profiles^35^, or perimortem factors^36^. Whatever the case, we found that homeostatic astrocytes constitute the majority of the astrocyte population, regardless of the severity of AD pathology, and express higher levels of typical homeostatic genes, including growth factor receptors, mitochondrial electron transport chain, and glutamate metabolism-related genes, whereas reactive astrocytes represent less than 25% of all astrocyte nuclei and express higher levels of aquaporins, heat shock proteins, metallothioneins, extracellular matrix, and cytoskeletal genes. Of note, many of these upregulated gene products have been demonstrated in AD reactive astrocytes via immunohistochemistry, especially near dense-core, typically neuritic, A*β* plaques^37^. Lastly, we also identify intermediate astrocytes, which constitute the second largest group after homeostatic astrocytes and are characterized by a gradient of transcriptomic changes between homeostatic and reactive astrocytes. Our trajectory analysis indicates that intermediate astrocytes represent transitional states between homeostatic and reactive astrocytes, rather than separate astrocyte subpopulations. This is consistent with a dynamic phenotypic change of astrocytes from homeostatic to reactive through these intermediate states in response to their microenvironment conditions, namely the presence of A*β* plaques and NFTs^12, 38–41^ and ongoing work will define the transcription factors involved in these transitions in AD^42^.

In summary, our five-region snRNA-seq study of astrocytes from 32 donors spanning the normal aging-AD continuum provides evidence of the regional diversity of astrocytes in the normal aging brain, demonstrates the complexity of their transcriptomic responses upon chronic exposure to ADNC, suggests A*β* plaque- and pTau-associated unique signatures of astrocytic reaction, and uncovers the existence of intermediate states that appear to be transitional between homeostatic and reactive astrocytes. Overall, our findings implicate astrocyte dysfunction in AD progression.

## METHODS

### Human specimens

Frozen (-80°C) brain specimens from 32 subjects were obtained from the Massachusetts Alzheimer’s Disease Research Center (MADRC) Brain Bank, including 11 Braak NFT 0/I/II (B1), five Braak NFT III/IV (B2), and 16 Braak NFT V/VI (B3). Donors or their next-of-kin provided written informed consent for the brain donation and the study was performed under the MADRC Neuropathology Core Brain Bank Institutional Review Board approval. Approximately 10-20mg of tissue from visual cortex was homogenized (Precellys CK14 beads), RNA was extracted (MagMAX mirVana Total RNA) and RNA Integrity Number (RIN) was measured on a Tapestation (Agilent) in order to select high-quality tissue for single nuclei RNA-seq. RIN value was measured from 130 donors, for which 83 met the selected cutoff of RIN ≥ 5. Of these 83 donors, 32 were selected based on neuropathological criteria and tissue availability. RIN values were additionally measured from the EC, BA20, BA46, V2 and V1 pieces used for snRNA-seq. Five brain regions were chosen based on Braak NFT staging: entorhinal cortex (EC), inferior temporal cortex (BA20), dorsolateral prefrontal cortex (BA46), visual association cortex (V2), and primary visual cortex (V1). Care was taken to dissect cortex from underlying white matter and include only cortex.

### Nuclei isolation and sorting

Nuclei were isolated and sorted following published procedures^8^. Briefly, 30-40 forty-μm-thick cryostat sections were lysed in a sucrose lysis buffer (10 mM Tris-HCl pH 8.0, 320 mM sucrose, 5 mM CaCl_2_, 3 μM Mg(Ac)_2_, 0.1 mM EDTA, 1 mM DTT, and 0.1% Triton X-100). The resulting lysates were filtered through a 70 μm-pore cell strainer and nuclei were purified from filtrates by ultracentrifugation (107,000xg for 1.5 h at 4°C) through a sucrose cushion (10 mM Tris-HCl pH 8.0, 1.8 M sucrose, 3 μM Mg(Ac)_2_, 0.1 mM EDTA, and 1 mM DTT). Supernatants were removed and pellets were resuspended in 2% BSA/PBS containing RNase inhibitor (0.2 U/μL, Roche). To enrich astrocyte nuclei, we followed a strategy of depletion of neurons and oligodendrocytes. Nuclei were labeled with AlexaFluor647-conjugated mouse monoclonal anti-NEUN antibody (clone 1B7, Novus Biologicals, NBP1-92693AF647) and PE-conjugated mouse monoclonal anti-OLIG2 antibody (clone 211F1.1, EMD Millipore, MABN50A4). After washes, nuclei were stained with Sytox blue (Thermo Fisher) and sorted on a BD FACSAria Fusion. For each sample, we collected Sytox^pos^NeuN^pos^Olig2^neg^ and Sytox^pos^NeuN^neg^Olig2^neg^.

### snRNA-seq library construction and sequencing

Single-nucleus cDNA libraries were constructed using the Chromium Single Cell 3‘ Reagents Kit V3 following the manufacturer’s instructions (10x Genomics). Samples were pooled and sequenced targeting at least 30k reads per cell on a HiSeq2000. Separation of NeuN^pos^Olig2^neg^ and NeuN^neg^Olig2^neg^ into unique libraries allowed for increased number of reads and detected genes in non-neuronal cells as compared to published studies containing non-enriched nuclei (Figure S1a),

### Phospho-tau and tau ELISA

Protein extracts were prepared from frozen tissue adjacent to that used for snRNA-seq by homogenizing in phosphate-buffered saline (PBS) and centrifuging at 3,000xg for 10 min at 4°C. The resulting supernatants were subjected to ELISA for total tau and tau phosphorylated at Threonin-231 using the MesoScale Discovery Phospho(Thr231)/Total Tau Kit (MSD, #K15121D) according to the manufacturer’s instructions. The plates were developed using MESO QuickPlex SQ120 Plate Reader (MSD). Total tau and phosphorylated tau concentrations were determined using the calibration curve.

### Immunohistochemistry

For A*β* immunohistochemistry, frozen cryostat sections adjacent to those used for snRNA-seq were subjected to immunohistochemistry with the mouse monoclonal anti N-terminal A*β* antibody clone 3D6. The staining was performed on a Leica BOND Rx automated stainer using DAB-based detection (Leica). Sections were scanned in a slide scanner (3DHistech, Panoramic 250) and area fraction (i.e., % area of tissue section occupied by 3D6-immunoreactive plaques) was measured using HALO software (Indica Labs, Albuquerque, NM, USA). For validation studies, 7-μm-thick formalin-fixed paraffin-embedded sections from the contralateral brain hemisphere of the same donors were dewaxed in xylenes, rehydrated in decreasing concentrations of ethanol, and subjected to antigen retrieval (microwaved in boiling citrate buffer 0.1 M, pH 6.0, with 0.05% Tween-20 at 95°C for 20 min), followed by blocking with 10% normal donkey serum in TBS for 1 h at room temperature. Primary antibodies (see Supplementary Table 8) were incubated in 5% normal donkey serum in TBS overnight at 4°C. Fluorescently-labeled secondary antibodies were incubated in 5% normal donkey serum in TBS for 2 h at room temperature. For Thioflavin-S staining, sections were incubated in thioflavin-S 0.05% dissolved in ethanol 50% for 8 min and washed with ethanol 80% and distilled water. Slides were coverslipped with DAPI-containing mounting media (Fluoromount-G DAPI, Southern Biotech) and scanned in a VS120 Olympus slide scanner with the same exposure time for each marker across brain regions (i.e., EC vs. V1) and/or diagnoses (i.e., control vs. AD). Immunohistochemistry images in figure panels have the same display settings across brain regions and diagnoses.

### *In situ* hybridization

To demonstrate *MAPT* expression in astrocytes, we performed 3-plex fluorescent *in situ* hybridization on cryostat sections from the temporal association cortex of selected control and AD donors using RNAScope technology (ACD, Biotechne), following manufacturer’s protocol. Probes to detect human *GFAP*, *MAPT*, and *SYP* (synaptophysin) mRNA (Cat# 311801-C2, 40899, and 311421-C3, respectively) were labeled with fluorescein, cyanine-3, and cyanine-5.5, respectively. Endogenous tissue autofluorescence was quenched with TrueBlack Lipofuscin Autofluorescence Quencher (Biotium Inc., USA). Sections were imaged at 63x magnification using an Olympus FV3000 confocal laser scanning microscope (Olympus, Tokyo, Japan).

### Bioinformatics analyses

#### snRNA-seq processing, cell type identification

Raw reads were processed using Cell Ranger 3.0.0 with default settings, pre-mRNA package and aligned to the human GRCh38 genome. For each brain region, we removed cells with fewer than 200 genes, greater than 20,000 unique molecular identifiers (UMIs), and/or greater than 20% mitochondrial genes, and used reciprocal principal component analysis (rPCA) integration based on the top 2,000 highly variable genes (HVG) to remove donor-specific effects using the Seurat^43^ R package (version 4.0.0). For the clustering of nuclei, integrated gene expression data were log-normalized, scaled, and subjected to PCA to choose the number of principal components for clustering, which was followed by non-linear dimensionality reduction via Uniform Manifold Approximation and Projection (UMAP) and visualized with UMAP plots. Next, nuclei from various cell types were annotated based on the relative expression levels of known genes. Astrocyte nuclei were identified by their high expression of *ADGRV1*, *ALDH1L1*, *AQP4*, and *GFAP*.

The following filters were applied to select the genes and astrocyte nuclei for downstream analyses: (1) 18,283 protein coding genes (i.e., non-coding genes were excluded); (2) genes with sum of counts >100 in at least 30% of samples (e.g., n=11,148 for BA20); (3) nuclei with <25,000 UMIs after gene filtering (to exclude multiplets) and >2,000 genes detected (i.e., with a count >0) (e.g., n=70,984 nuclei for BA20). After filtering, a second round of integration was performed via canonical correlation analysis (CCA) to regress out donor-specific effects as well as the percentage of mitochondrial genes. Briefly, data were scaled, PCA was applied on scaled data, and the number of principal components used in clustering was decided based on the Elbow plot. Finally, the clustering was conducted at a 0.3 resolution for all brain regions, non-linear dimensionality reduction was performed via UMAP, and results were visualized with UMAP plots. Small donor-specific clusters and clusters contaminated with other cell types were removed prior to downstream analyses. Details are described in Supplementary Table 9.

#### Analysis of normal aging brains

Nuclei from control brains were integrated by rPCA and clustered using FindClusters at 0.4 resolution. Markers for each cluster were identified using FindAllMarkers and functional enrichment of clusters was performed against the msigDB database^44^ using a hypergeometric test. Differential expression of each brain region relative to other regions was also performed using FindAllMarkers. Differentially expressed genes (DEGs) that pass the average fold-change (FC) cut off (FC > 1.2 for up-regulated genes and FC < 0.8 for the down-regulated genes), had a false discovery rate (FDR) < 0.05 and were expressed in greater than 50% of the cells in the given brain region are reported. To find association of gene expression with pTau pathology, we used MAST^45^ to perform zero-inflated regression analysis by fitting a mixed model. The model specification was:

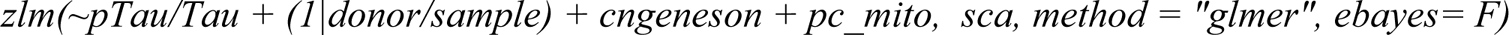

where *cngeneson* is the cellular detection rate and *pc_mito* is the percentage of counts mapping to mitochondrial genes.

#### Identification of gene sets following distinct spatial and temporal trajectories

To identify gene sets that followed distinct spatial trajectories along the neural network, we first performed a differential expression analysis comparing each node of the network with the next (i.e., EC vs. ITG, ITG vs. PFC, PFC vs. V2, and V2 vs. V1) using FindMarkers() with DonorID and Pathology Stage as latent variables. We selected DEGs that were logFC > 0.25, adjusted *p* < 0.05 and percentage expression greater than 10% in both regions. Next, for describing the spatial gene patterns, we calculated the standardized gene expression score scaling across brain regions for each donor and then averaged those across all donors for each brain region. The resulting mean standardized gene expression scores were grouped into six gene sets by spectral clustering using SNFTool (v.2.3.1)^46^ with k = 6. To find the overlap with the EC/V1 high or low genes, we used a hypergeometric test. And, to compute the association of the gene sets with A*β* and pTau/Tau local measures, we used a mixed effects model:

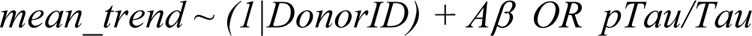

For the temporal gene patterns, we similarly performed a differential expression analysis comparing two “adjacent” pathology stages (Stage 2 *vs*. Stage 1, Stage 3 *vs*. Stage 2, Stage 4 *vs*. Stage 3) using FindMarkers() with Region as latent variable. We selected DEGs that were logFC > 0.25, adjusted *p* < 0.05 and percentage expression greater than 10% in both regions. We then computed the standardized gene expression scores by scaling across all the samples for a gene and then computed the average standardized gene expression score for each pathology stage. The resulting scores were similarly used as input for the spectral clustering using SNFTool (v.2.3.1)^46^ with k = 6.

#### Differential Expression (DE) analysis and functional enrichment of astrocyte subclusters

To characterize the transcriptomic signatures defining distinct astrocyte subclusters, DE analyses were conducted comparing the expression level of each gene in each astrocyte subcluster with that of each of the other astrocyte subclusters detected within the same brain region via logistic regression using the FindAllMarkers function within the package Seurat, with no filters and regressing out donor-specific effects (i.e., latent variable = donor ID). To map astrocyte subclusters across brain regions, the log fold-change of each astrocyte subcluster relative to all other subclusters in that brain region was correlated to that of all astrocyte subclusters from all other brain regions via Spearman’s correlation. The union of highly variable genes in each dataset was used for this task. The Spearman’s correlation coefficients were visualized with a heatmap.

To characterize the functional alteration defining distinct astrocyte subclusters, we performed hypergeometric test against the Reactome database for each astrocyte subclusters using MSigDB^44^. The up-regulated genes for each astrocytes subclusters were defined with FC > 1.2, adjusted *p* < 0.05 and percentage expression greater than 40% in given cluster. The down-regulated genes for each astrocyte subcluster were defined with FC < 0.8, adjusted *p* < 0.05 and percentage expression greater than 40% in given cluster. The resulting enriched pathways were clustered using a Jaccard index of similarly in gene membership; representative pathways were selected by expert and illustrated using *ggplot* in R and the full list of significant pathways were reported in supplementary tables.

#### Trajectory Analysis

Astrocytes were sampled equally from each cluster (n = 1,000 for each cluster, total n = 10,000). The sampled data was normalized, and log transformed in Python via *scanpy* (v.1.7.1)^47^. The top 1,500 highly variable genes were selected with flavor = ‘cell_ranger’ for the principal component analysis. Then, diffusion maps were estimated via *palantir.utils.run_diffusion_maps* (Palantir v1.0.0, harmony v0.1.4) as a low dimensional embedding of the data^48^. As recommended by the author of the Palantir^48^, we computed force-directed layouts by the same adaptative kernel that was used to estimate diffusion maps for visualization of the trajectory. Markov affinity-based graph imputation of cells (MAGIC) imputation method developed by Pe’er lab was performed by *palantir.utils.run_magic_imputation* for imputation^49^. After above processing steps, we did trajectory analysis from astH1 to astR1 with predefined start and terminal cells. According to the force-directed layouts visualization, the astH1 cell with min force-directed layouts and closest to the diagonal was selected as start cell, and the astR1 cell with max force-directed layouts and closest to the diagonal was defined as terminal cell. Finally, the trends of 1,500 highly variable genes along the trajectory were clustered by *palantir.presults.compute_gene_trends*. All analyses were performed with Python 3.8.5.

## DATA AVAILABILITY

The snRNA-seq dataset generated for the study will be made available at GEO and SRA (accession #xxxxxxxx) upon acceptance of the paper. Genes and pathways information are available as Supplementary Tables. Data can also be queried via the interactive web application at https://alzdatalens.partners.org/?page=ad-progression-study. The processed single-cell gene expression data and metadata can also be downloaded directly from the website.

## CODE AVAILABILITY

All code generated during this study is accessible at https://github.com/mindds/ad-progression-study.

## AUTHOR CONTRIBUTIONS

AS-P, BTH, REB, and SD conceived the study and designed experiments, and together with ZL, designed the bioinformatics analyses. MEW led the protocol development for single-nucleus sequencing from the human brain. TP performed nuclei isolation, MEW performed fluorescence-activated nuclei sorting (FANS), JT and AA generated the snRNA-Sequencing libraries, KZ and FL quantified the A*β* plaque burden, MH performed the pTau/Tau ELISAs, CM-C performed the fluorescent immunohistochemistry validation, and LAW performed the RNAScope validation. RJ, AB, AW, TK, and LG contributed to the analyses and AN developed the shiny web interface. BTH, KB, RVT, and EK acquired funding and initiated the study. AS-P and SD wrote the manuscript; MEW, REB, and ZL contributed to the Methods. All authors approved of, and contributed to, the final version of the manuscript.

## ACKNOWLEDGEMENTS

We would like to thank the patients and families involved in research at the Massachusetts Alzheimer’s disease Research Center. We also thank Patrick Dooley and Tessa Connors from the MADRC brain bank. We acknowledge funding from the following sources: NIH/NIA (P30AG062421 to BTH and SD, K08AG034069 to AS-P), Alzheimer’s Association (AACF-17-524184 and AACF-17-524184-RAPID to AS-P), and Real Colegio Complutense at Harvard University (Research Fellowship to CM-C).

---

**Figure S1.**
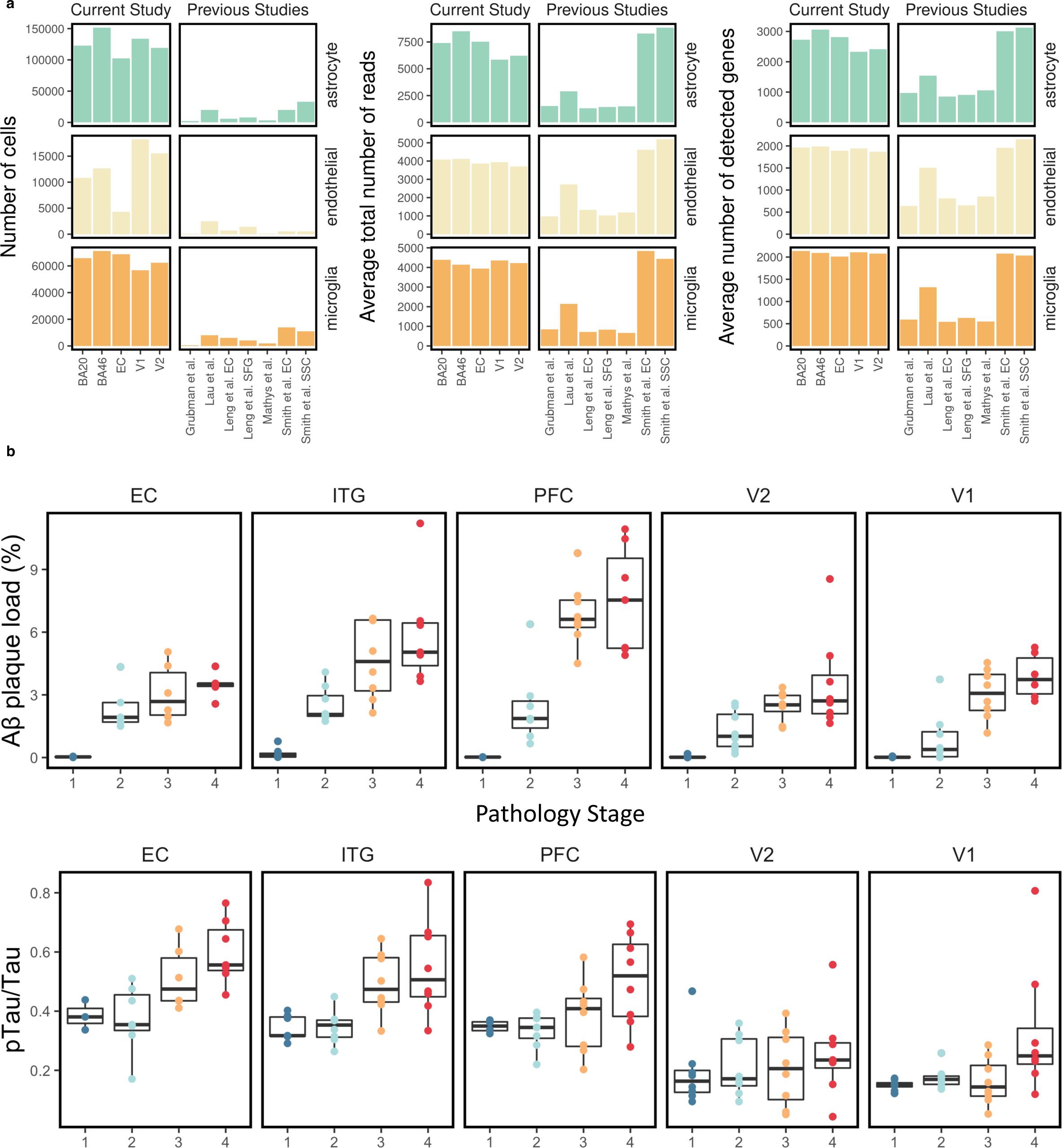
Characteristics of our snRNA-seq study. **a**, Comparison of our AD progression study data with previously published snRNA-seq AD datasets in number of cells, average total number of reads per cell, and average number of detected genes (counts > 0) per cell. **b**, Box plots show the Aβ plaque load (immunoreactive % area fraction) and pTau/Tau ratio (measured by ELISA) of the 32 donors across pathology stages and brain regions.

**Figure S2.**
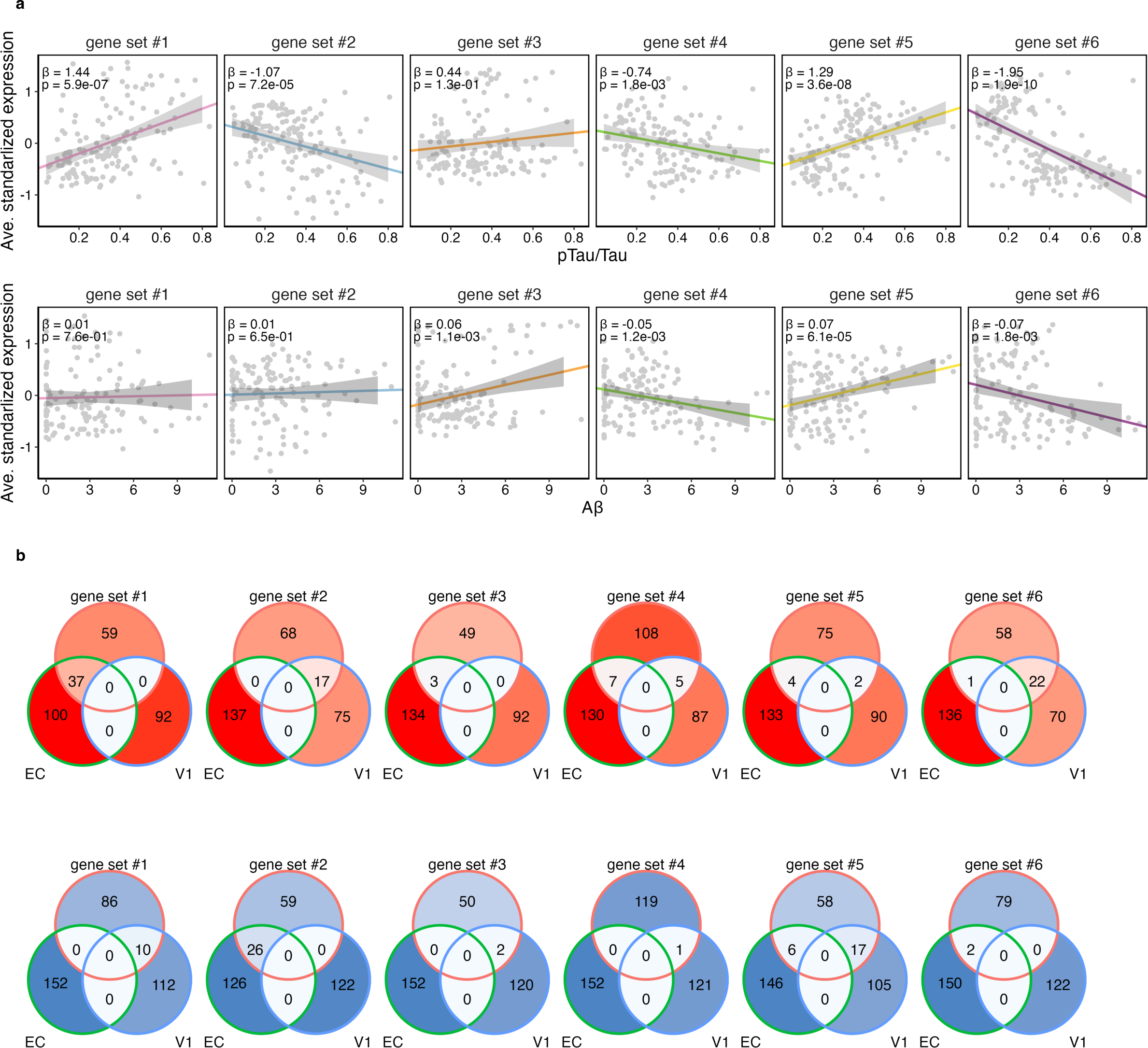
Astrocyte transcriptomic changes along the stereotypical spatial progression of AD. **a**, Scatter dot plots show the correlation between the average standardized expression of each spatial trajectory gene set in each donor and their Aβ plaque load (top) or pTau/Tau ratio (bottom). **b**, Venn diagrams show the overlap between each of the spatial trajectory gene sets and the EC- and V1-upregulated (red) and -downregulated (blue) genes.

**Figure S3.**
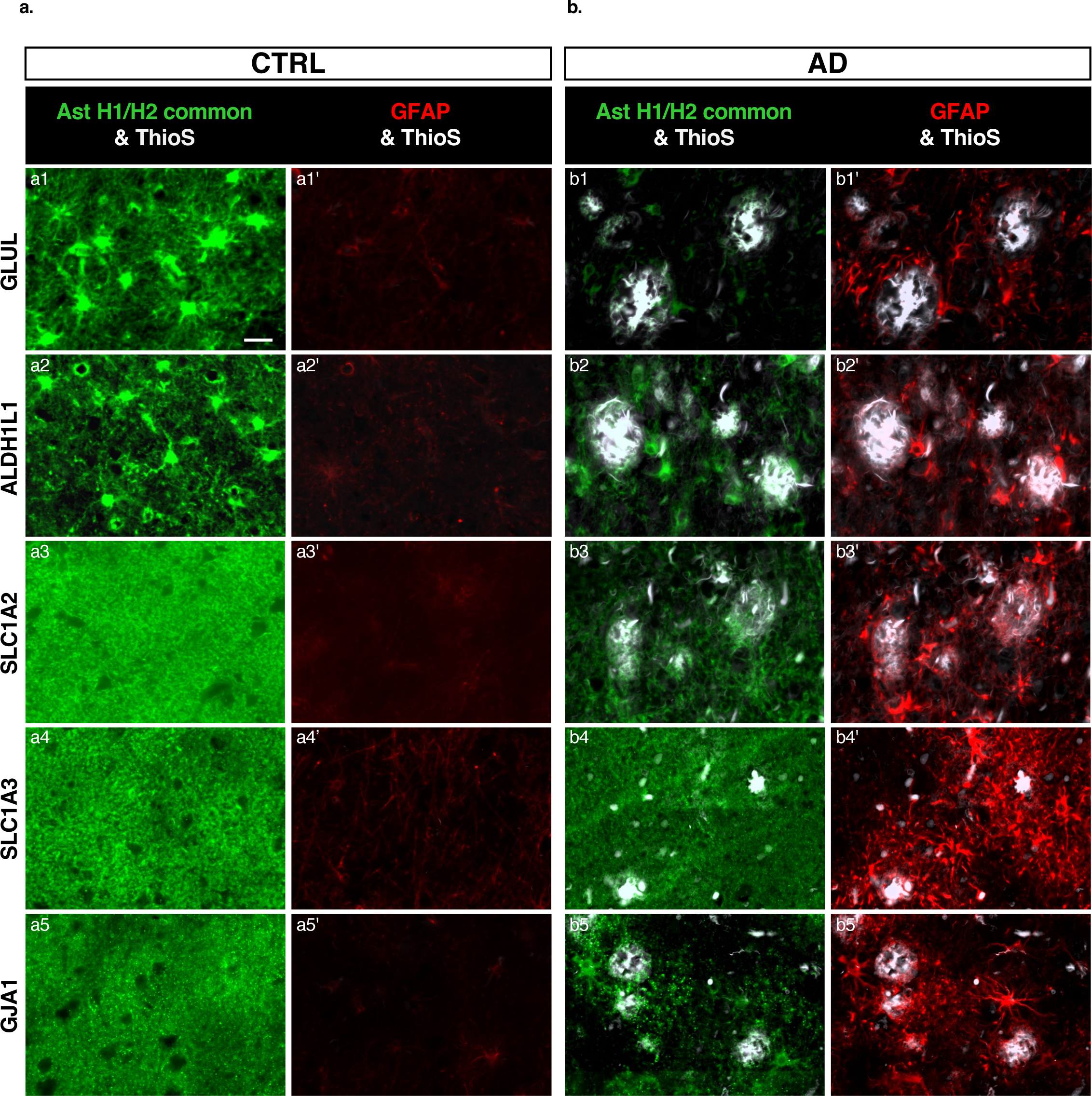
Immunohistochemical analysis of homeostatic astrocyte markers. Double fluorescent immunohistochemistry for GFAP and relevant homeostatic astrocyte markers with Thioflavin-S (ThioS) counterstaining in control (CTRL, **a**) and AD (**b**) formalin-fixed paraffin-embedded sections from the temporal association cortex. Note some reduced expression of these markers in reactive (GFAP+) astrocytes surrounding dense-core (ThioS+) Aβ plaques in AD but an apparent redistribution of GJA1 towards Aβ plaques. Note: photomicrographs from CTRL and AD donors were taken with similar exposure time and display settings for appropriate comparison. Scale bar: 20 μm.

**Figure S4.**
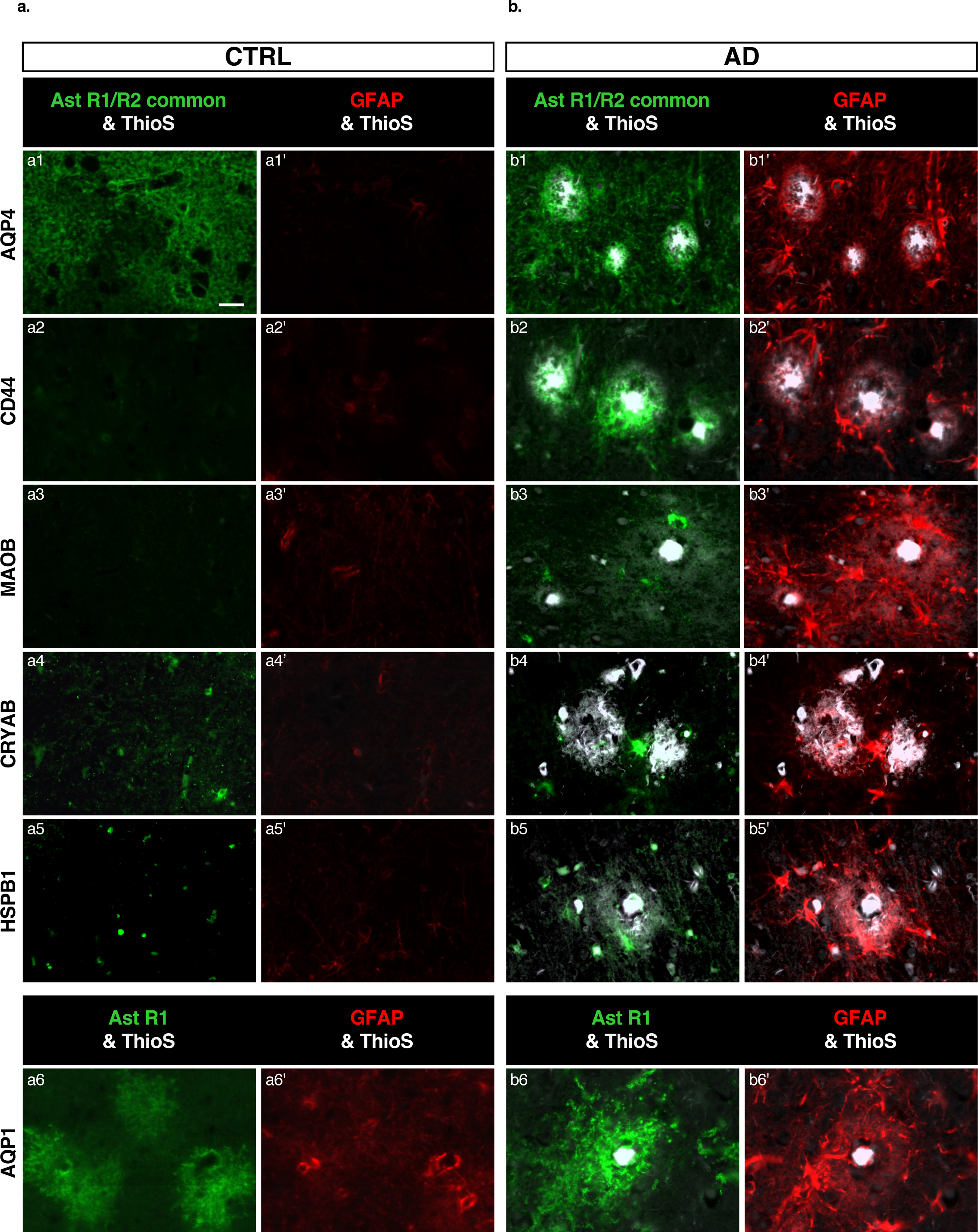
Immunohistochemical analysis of reactive astrocyte markers. Double fluorescent immunohistochemistry for GFAP and relevant reactive astrocyte markers with Thioflavin-S (ThioS) counterstaining in control (CTRL, **a**) and AD (**b**) formalin-fixed paraffin-embedded sections from the temporal association cortex. Note the increased expression of these markers in reactive (GFAP+) astrocytes surrounding dense-core (ThioS+) Aβ plaques and an apparent redistribution of AQP4 towards Aβ plaques in AD. Note: photomicrographs from CTRL and AD donors were taken with similar exposure time and display settings for appropriate comparison. Scale bar: 20 μm.

**Figure S5.**
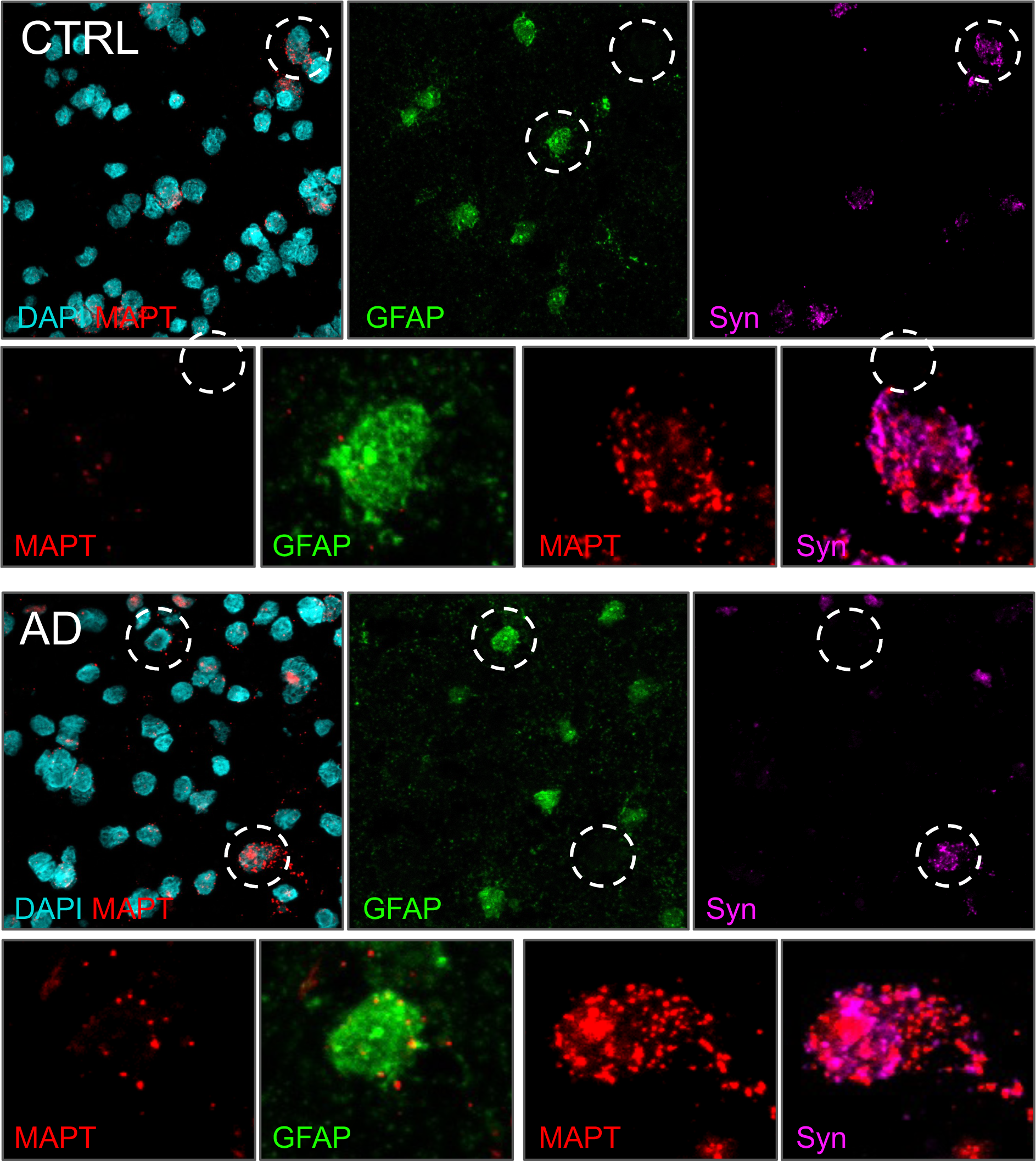
*In situ* hybridization (RNAScope) analysis of *MAPT*. RNAScope for *MAPT*, *GFAP*, and *SYP* (encoding the neuronal synaptic marker synaptophysin) reveals some expression of *MAPT* in astrocytes.

## REFERENCES

1. Verkhratsky, A. & Nedergaard, M. Physiology of Astroglia. Physiol Rev 98, 239–389, doi:10.1152/physrev.00042.2016 (2018).

2. Escartin, C. et al. Reactive astrocyte nomenclature, definitions, and future directions. Nat Neurosci 24, 312–325, doi:10.1038/s41593-020-00783-4 (2021).

3. Mathys, H. et al. Single-cell transcriptomic analysis of Alzheimer’s disease. Nature, doi:10.1038/s41586-019-1195-2 (2019).

4. Grubman, A. et al. A single-cell atlas of entorhinal cortex from individuals with Alzheimer’s disease reveals cell-type-specific gene expression regulation. Nature Neuroscience, doi:10.1038/s41593-019-0539-4 (2019).

5. Leng, K. et al. Molecular characterization of selectively vulnerable neurons in Alzheimer’s disease. Nature Neuroscience 24, 276–287, doi:10.1038/s41593-020-00764-7 (2021).

6. Lau, S.-F., Cao, H., Fu, A. K. Y. & Ip, N. Y. Single-nucleus transcriptome analysis reveals dysregulation of angiogenic endothelial cells and neuroprotective glia in Alzheimer’s disease. Proceedings of the National Academy of Sciences 117, 25800–25809, doi:10.1073/pnas.2008762117 (2020).

7. Sadick, J. S. et al. Astrocytes and oligodendrocytes undergo subtype-specific transcriptional changes in Alzheimer’s disease. Neuron 110, 1788–1805 e1710, doi:10.1016/j.neuron.2022.03.008 (2022).

8. Gerrits, E. et al. Distinct amyloid-β and tau-associated microglia profiles in Alzheimer’s disease. Acta Neuropathologica 141, 681–696, doi:10.1007/s00401-021-02263-w (2021).

9. Smith, A. M. et al. Diverse human astrocyte and microglial transcriptional responses to Alzheimer’s pathology. Acta Neuropathologica, doi:10.1007/s00401-021-02372-6 (2021).

10. Braak, H. & Braak, E. Neuropathological stageing of Alzheimer-related changes. Acta Neuropathol 82, 239–259, doi:10.1007/BF00308809 (1991).

11. Braak, H., Alafuzoff, I., Arzberger, T., Kretzschmar, H. & Del Tredici, K. Staging of Alzheimer disease-associated neurofibrillary pathology using paraffin sections and immunocytochemistry. Acta Neuropathol 112, 389–404, doi:10.1007/s00401-006-0127-z (2006).

12. Serrano-Pozo, A. et al. Reactive glia not only associates with plaques but also parallels tangles in Alzheimer’s disease. Am J Pathol 179, 1373–1384, doi:10.1016/j.ajpath.2011.05.047 (2011).

13. Perez-Nievas, B. G. & Serrano-Pozo, A. Deciphering the Astrocyte Reaction in Alzheimer’s Disease. Front Aging Neurosci 10, 114, doi:10.3389/fnagi.2018.00114 (2018).

14. Kunkle, B. W. et al. Genetic meta-analysis of diagnosed Alzheimer’s disease identifies new risk loci and implicates Abeta, tau, immunity and lipid processing. Nat Genet 51, 414–430, doi:10.1038/s41588-019-0358-2 (2019).

15. Barker, S. J. et al. MEF2 is a key regulator of cognitive potential and confers resilience to neurodegeneration. Sci Transl Med 13, eabd7695, doi:10.1126/scitranslmed.abd7695 (2021).

16. Van Hoesen, G. W., Hyman, B. T. & Damasio, A. R. Entorhinal cortex pathology in Alzheimer’s disease. Hippocampus 1, 1–8, doi:10.1002/hipo.450010102 (1991).

17. Kobayashi, E. et al. Activated forms of astrocytes with higher GLT-1 expression are associated with cognitive normal subjects with Alzheimer pathology in human brain. Sci Rep 8, 1712, doi:10.1038/s41598-018-19442-7 (2018).

18. Ortinski, P. I. et al. Selective induction of astrocytic gliosis generates deficits in neuronal inhibition. Nat Neurosci 13, 584–591, doi:10.1038/nn.2535 (2010).

19. Funkhouser, E. B. The Visual Cortex, Its Localization, Histological Structure, and Physiological Function. J Exp Med 21, 617–628, doi:10.1084/jem.21.6.617 (1915).

20. Kohler, S., Winkler, U. & Hirrlinger, J. Heterogeneity of Astrocytes in Grey and White Matter. Neurochem Res 46, 3–14, doi:10.1007/s11064-019-02926-x (2021).

21. Allen, D. E. et al. Fate mapping of neural stem cell niches reveals distinct origins of human cortical astrocytes. Science 376, 1441–1446, doi:10.1126/science.abm5224 (2022).

22. Montal, V. et al. Network Tau spreading is vulnerable to the expression gradients of APOE and glutamatergic-related genes. Sci Transl Med 14, eabn7273, doi:10.1126/scitranslmed.abn7273 (2022).

23. Jiwaji, Z. et al. Reactive astrocytes acquire neuroprotective as well as deleterious signatures in response to Tau and Ass pathology. Nat Commun 13, 135, doi:10.1038/s41467-021-27702-w (2022).

24. Galea, E. et al. Multi-transcriptomic analysis points to early organelle dysfunction in human astrocytes in Alzheimer’s disease. Neurobiol Dis 166, 105655, doi:10.1016/j.nbd.2022.105655 (2022).

25. Caglayan, E., Liu, Y. & Konopka, G. Neuronal ambient RNA contamination causes misinterpreted and masked cell types in brain single-nuclei datasets. Neuron, doi:10.1016/j.neuron.2022.09.010 (2022).

26. Zimmer, E. R. et al. [(18)F]FDG PET signal is driven by astroglial glutamate transport. Nat Neurosci 20, 393–395, doi:10.1038/nn.4492 (2017).

27. Salvado, G. et al. Reactive astrogliosis is associated with higher cerebral glucose consumption in the early Alzheimer’s continuum. Eur J Nucl Med Mol Imaging, doi:10.1007/s00259-022-05897-4 (2022).

28. March-Diaz, R. et al. Hypoxia compromises the mitochondrial metabolism of Alzheimer’s disease microglia via HIF1. Nature Aging 1, 385–399, doi:10.1038/s43587-021-00054-2 (2021).

29. Lananna, B. V. et al. Chi3l1/YKL-40 is controlled by the astrocyte circadian clock and regulates neuroinflammation and Alzheimer’s disease pathogenesis. Sci Transl Med 12, doi:10.1126/scitranslmed.aax3519 (2020).

30. Serrano-Pozo, A., Das, S. & Hyman, B. T. APOE and Alzheimer’s disease: advances in genetics, pathophysiology, and therapeutic approaches. Lancet Neurol 20, 68–80, doi:10.1016/S1474-4422(20)30412-9 (2021).

31. Guttenplan, K. A. et al. Neurotoxic reactive astrocytes induce cell death via saturated lipids. Nature 599, 102–107, doi:10.1038/s41586-021-03960-y (2021).

32. La Joie, R. et al. Prospective longitudinal atrophy in Alzheimer’s disease correlates with the intensity and topography of baseline tau-PET. Sci Transl Med 12, doi:10.1126/scitranslmed.aau5732 (2020).

33. Zhou, Y. et al. Human and mouse single-nucleus transcriptomics reveal TREM2-dependent and TREM2-independent cellular responses in Alzheimer’s disease. Nature Medicine 26, 131–142, doi:10.1038/s41591-019-0695-9 (2020).

34. Wang, M. et al. Guidelines for bioinformatics of single-cell sequencing data analysis in Alzheimer’s disease: review, recommendation, implementation and application. Mol Neurodegener 17, 17, doi:10.1186/s13024-022-00517-z (2022).

35. Thrupp, N. et al. Single-Nucleus RNA-Seq Is Not Suitable for Detection of Microglial Activation Genes in Humans. Cell Rep 32, 108189, doi:10.1016/j.celrep.2020.108189 (2020).

36. Marsh, S. E. et al. Dissection of artifactual and confounding glial signatures by single-cell sequencing of mouse and human brain. Nat Neurosci 25, 306–316, doi:10.1038/s41593-022-01022-8 (2022).

37. Viejo, L. et al. Systematic review of human post-mortem immunohistochemical studies and bioinformatics analyses unveil the complexity of astrocyte reaction in Alzheimer’s disease. Neuropathol Appl Neurobiol 48, e12753, doi:10.1111/nan.12753 (2022).

38. Serrano-Pozo, A., Gomez-Isla, T., Growdon, J. H., Frosch, M. P. & Hyman, B. T. A phenotypic change but not proliferation underlies glial responses in Alzheimer disease. Am J Pathol 182, 2332–2344, doi:10.1016/j.ajpath.2013.02.031 (2013).

39. Serrano-Pozo, A. et al. Differential relationships of reactive astrocytes and microglia to fibrillar amyloid deposits in Alzheimer disease. J Neuropathol Exp Neurol 72, 462–471, doi:10.1097/NEN.0b013e3182933788 (2013).

40. Serrano-Pozo, A., Betensky, R. A., Frosch, M. P. & Hyman, B. T. Plaque-Associated Local Toxicity Increases over the Clinical Course of Alzheimer Disease. Am J Pathol 186, 375–384, doi:10.1016/j.ajpath.2015.10.010 (2016).

41. Munoz-Castro, C. et al. Cyclic multiplex fluorescent immunohistochemistry and machine learning reveal distinct states of astrocytes and microglia in normal aging and Alzheimer’s disease. J Neuroinflammation 19, 30, doi:10.1186/s12974-022-02383-4 (2022).

42. Burda, J. E. et al. Divergent transcriptional regulation of astrocyte reactivity across disorders. Nature 606, 557–564, doi:10.1038/s41586-022-04739-5 (2022).

43. Hao, Y. et al. Integrated analysis of multimodal single-cell data. Cell 184, 3573–3587 e3529, doi:10.1016/j.cell.2021.04.048 (2021).

44. Liberzon, A. et al. Molecular signatures database (MSigDB) 3.0. Bioinformatics 27, 1739–1740, doi:10.1093/bioinformatics/btr260 (2011).

45. Finak, G. et al. MAST: a flexible statistical framework for assessing transcriptional changes and characterizing heterogeneity in single-cell RNA sequencing data. Genome Biol 16, 278, doi:10.1186/s13059-015-0844-5 (2015).

46. Wang, B. et al. Similarity network fusion for aggregating data types on a genomic scale. Nat Methods 11, 333–337, doi:10.1038/nmeth.2810 (2014).

47. Wolf, F. A., Angerer, P. & Theis, F. J. SCANPY: large-scale single-cell gene expression data analysis. Genome Biol 19, 15, doi:10.1186/s13059-017-1382-0 (2018).

48. Setty, M. et al. Characterization of cell fate probabilities in single-cell data with Palantir. Nat Biotechnol 37, 451–460, doi:10.1038/s41587-019-0068-4 (2019).

49. van Dijk, D. et al. Recovering Gene Interactions from Single-Cell Data Using Data Diffusion. Cell 174, 716–729 e727, doi:10.1016/j.cell.2018.05.061 (2018).

